# AzuCR RNA modulates carbon metabolism as a dual-function RNA in *Escherichia coli*

**DOI:** 10.1101/2021.04.27.441574

**Authors:** Medha Raina, Jordan J. Aoyama, Shantanu Bhatt, Brian J. Paul, Gisela Storz

## Abstract

Bacteria have evolved small RNAs (sRNAs) to regulate numerous biological processes and stress responses. While sRNAs generally are considered to be “noncoding”, a few have been found to also encode a small protein. Here we describe one such dual-function RNA that modulates carbon utilization in *Escherichia coli*. The 164 nucleotide RNA was previously shown to encode a 28 amino acid protein (denoted AzuC). We discovered the membrane-associated AzuC protein interacts with GlpD, the aerobic glycerol-3-phosphate dehydrogenase, leading to increased GlpD activity. Overexpression of the RNA encoding AzuC results in a growth defect in glycerol and galactose medium. The defect in galactose medium was still observed for a stop codon mutant derivative, suggesting a potential regulatory role for the RNA. Consistent with this observation, we found that *cadA* and *galE* are repressed by base pairing with the RNA (denoted AzuCR). Interestingly, translation of AzuC interferes with the observed repression of *cadA* and *galE* by AzuCR and base pairing interferes with AzuC translation, demonstrating that the translation and base pairing functions compete.

## Introduction

Bacteria are exposed to rapidly changing environmental conditions including variations in carbon availability, pH, temperature, and osmolarity, to name a few. To survive these fluctuating conditions, bacterial cells have fast, flexible, and energy-efficient mechanisms to regulate protein levels and activity. Changes in nutrient availability or stress detected by the cell are transduced into changes in the activation or repression of transcription, post-transcriptional changes to mRNA stability, modulation of mRNA translation as well as the modification of protein stability or activity.

The cAMP receptor protein (CRP), a sequence-specific DNA binding protein, is a key regulator of transcription in response to changes in carbon source availability in *E. coli*. When the levels of the preferred carbon source glucose are low, the small molecule cAMP is synthesized. cAMP binds and activates the highly-conserved CRP transcription factor, which in turn activates genes for the uptake and utilization of alternative carbon sources in a process termed carbon catabolite repression (CCR) (reviewed in (Soberón-Chávez *et al*., 2017)).

Several small RNAs (sRNAs), which modulate the stability or translation of mRNAs through short base pairing interactions and are major post-transcriptional regulators in bacteria (reviewed in (Wagner & Romby, 2015)), also have been found to impact carbon metabolism in *E. coli*. These include GlmZ, ChiX, SgrS and the CRP-repressed sRNAs Spot 42 and CyaR (reviewed in (Papenfort & Vogel, 2014, Durica-Mitic *et al*., 2018)). The base-pairing sRNAs require RNA chaperones such as Hfq and ProQ for their stability and optimal pairing with their target mRNAs (reviewed in (Holmqvist & Vogel, 2018, Updegrove *et al*., 2016, Olejniczak & Storz, 2017)).

Base-pairing sRNAs generally are thought not to encode proteins and thus are often referred to as noncoding RNAs. However, a few sRNAs have been shown to be translated to produce small proteins and thus are denoted “dual-function RNAs” (reviewed in (Raina *et al*., 2018)). Computational analyses of the genomes of fourteen phylogenetically diverse bacteria predicted that a number of other sRNAs contain small open reading frames (sORFs) that could encode proteins between 10-50 amino acids (Friedman *et al*., 2017). Nevertheless, translation of these sORFs has only been documented in a limited number of cases. Even fewer sRNA-encoded protein products have experimentally been demonstrated to have a function.

Small proteins of less than 50 amino acids generally have been long overlooked due to many challenges related to their annotation and biochemical detection. The few that have now been studied show that small proteins modulate diverse cellular functions ranging from morphogenesis and cell division to transport, enzymatic activities, regulatory networks, and stress responses by forming complexes with larger proteins (reviewed in (Storz *et al*., 2014, Hemm *et al*., 2020)).

To date, the only dual-function RNA characterized in *E. coli* is SgrS (Wadler & Vanderpool, 2007). The SgrS RNA was first found to protect cells against elevated levels of glucose phosphate by regulating the stability and translation of mRNAs encoding proteins involved in glucose transport and catabolism (Vanderpool & Gottesman, 2004). The RNA subsequently was shown to encode a 43 amino acid protein, SgrT, which interacts with the glucose importer PtsG to block glucose transport and promote utilization of nonpreferred carbon sources to maintain growth during glucose-phosphate stress (Lloyd *et al*., 2017, Wadler & Vanderpool, 2007). Thus both the sRNA and its encoded small protein act together to repress glucose import to relieve glucose phosphate stress.

The 164 nt RNA initially denoted IS092 or IsrB (now denoted AzuCR) was identified in a bioinformatic search to find sRNA genes in *E. coli* (Chen *et al*., 2002), but not characterized as an sRNA. Later, this RNA was shown to encode an 28 amino acid sORF (Hemm *et al*., 2008) (Fig 1A). Synthesis of the small protein was documented by the detection of a tagged derivative (Hemm *et al*., 2008) and is supported by data indicating ribosome binding to the RNA (Weaver *et al*., 2019) (Fig EV1A). While the protein, denoted AzuC, is only conserved in a limited number of enteric bacteria (Fig EV1B), expression of AzuC was found to be highly regulated. The levels of the tagged small protein were elevated for growth in glucose compared to glycerol due to CRP-mediated repression in the absence of glucose (Hemm *et al*., 2010). AzuC-SPA levels also were shown to be reduced under anaerobic conditions but induced upon exposure to low pH, high temperature, and hydrogen peroxide suggesting an important role in cellular stress responses (Hemm *et al*., 2010).

**Figure 1.**
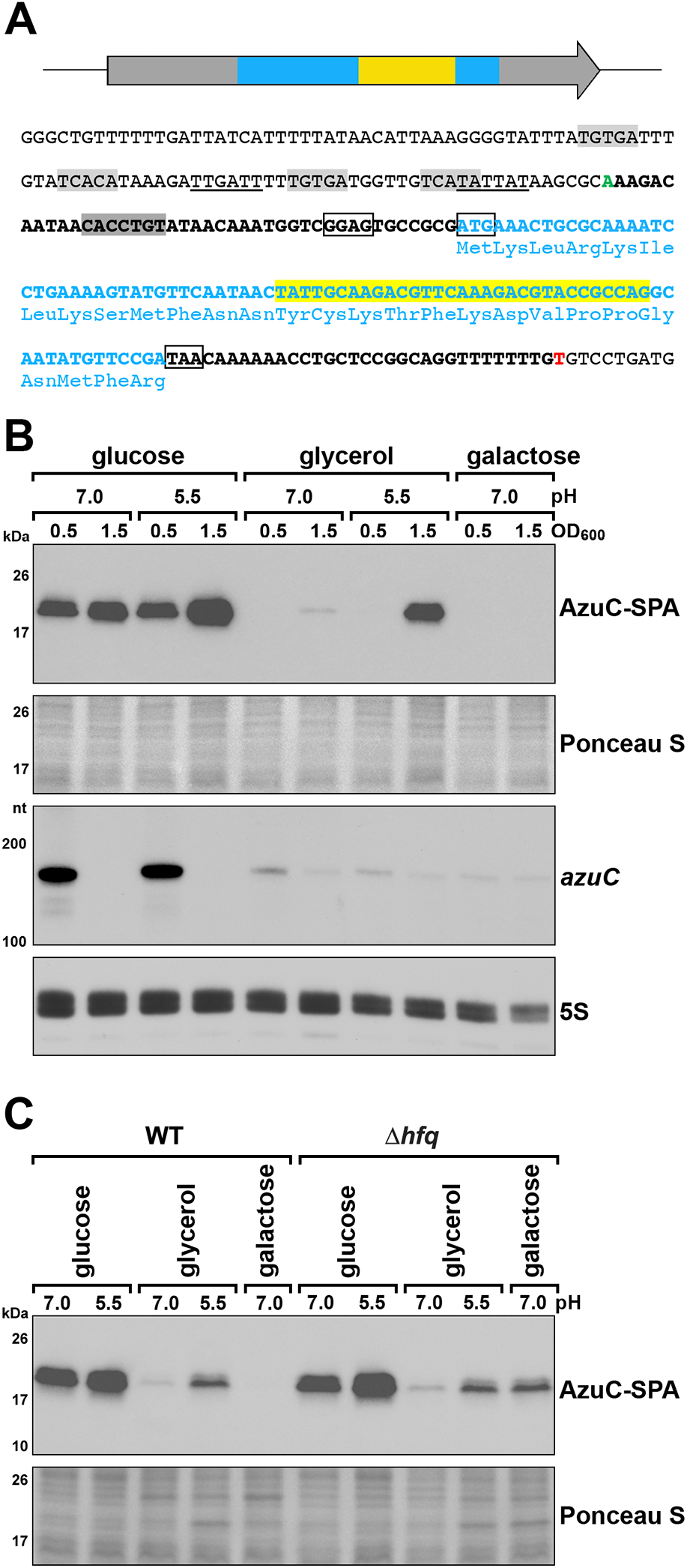
AzuC protein synthesis is regulated at a post-transcriptional level. A Diagram of the AzuCR RNA and sequence of the *azuC* promoter and coding region. Boxes and text in light blue denote AzuC coding sequence and yellow box and highlighted text denote region of base pairing with target mRNAs. The AzuCR transcript is indicated in bold with the +1 site of transcription (position 1988001 of the *E. coli* K-12 genome) in green font and the 3’ end of the transcript in red font. The ribosome binding site and the start and stop codons of the AzuC ORF are indicated by black boxes. Potential σ^70^ −10 and −35 sequences are underlined, the predicted CRP binding sites are highlighted in light gray (Hemm *et al*., 2010), and the region targeted by the FnrS sRNA is highlight in dark gray. B Immunoblot blot analysis of AzuC-SPA levels (top) and northern blot analysis of *azuC* mRNA levels (bottom) for cells grown in with different carbon sources. Cultures of the *azuC-SPA::kan* (GSO351) or unmarked (MG1655) strains were grown in M63 medium supplemented with glucose, glycerol, or galactose at pH 7.0 or 5.5. Samples were taken at OD_600_ ∼0.5 and 1.5. *α*-FLAG antibody was used to detect the SPA tag. The membrane was stained with Ponceau S stain to control for loading. The *azuC* mRNA and 5S RNA were detected by oligonucleotide probes specific to each of these transcripts. C Immunoblot blot analysis of AzuC-SPA levels in *hfq^+^* (GSO351) and Δ*hfq* (GSO1007) cells grown in with different carbon sources. Strains were grown in the same media as in (B) and collected at OD_600_ ∼1.5. Immunoblot blot analysis also was carried out as in (B).

Here we show that AzuC is associated with the membrane and binds GlpD, an enzyme required for glycerol catabolism, increasing GlpD activity. Additionally, we document that the transcript acts as a regulatory sRNA, denoted AzuR, repressing expression of *cadA*, a lysine decarboxylase involved in maintaining pH homeostasis, and *galE*, encoding UDP-glucose 4-epimerase, through direct base pairing. Thus, AzuCR has mRNA and sRNA activities in two different pathways. The protein coding and base-pairing sequences overlap, and we find that there is inherent competition between the two activities. Intriguingly, while the transcript base pairs with other mRNAs as a regulator, translation of AzuC itself is repressed by the FnrS sRNA, an sRNA that also represses GlpD synthesis.

## Results

### AzuC protein and mRNA levels are discordant for cells grown in glucose and low pH glycerol

Previous analysis of chromosomally-encoded AzuC, which was C-terminally tagged with the sequential peptide affinity (SPA) tag, showed that AzuC-SPA protein levels were elevated in cells grown in minimal medium with glucose compared to glycerol as well as in pH 5.5 compared to pH 7.5, and decreased under anerobic conditions (Hemm *et al*., 2010). The decreased levels in minimal glycerol medium and part of the pH-induction were attributed to CRP-mediated repression of *azuC* mRNA transcription.

To further evaluate the conditions under which AzuC-SPA and *azuC* mRNAs levels are highest, strains were cultured in M63 media supplemented with glucose or glycerol at pH 7.0 and 5.5, and in M63 galactose at pH 7.0 (the strain was unable to grow in M63 galactose at pH 5.5). Cells were collected in exponential (OD_600_ ∼0.5) and stationary (OD_600_ ∼1.5) phase (Fig 1B). As observed previously, AzuC-SPA levels were significantly higher in glucose compared to glycerol and galactose. A notable exception was the elevated AzuC-SPA levels in cells grown to stationary phase in glycerol at pH 5.5. As expected for a CRP-regulated transcript, *azuC* mRNA levels were low for all conditions except for cells growing exponentially in glucose. The discordance between AzuC-SPA protein levels and *azuC* mRNA levels in glycerol at pH 5.5 raised the possibility that translation and/or RNA or protein stability is regulated and that the protein and RNA may have different roles.

Hfq is a key regulator of posttranscriptional regulation in many bacterial cells (reviewed in (Holmqvist & Vogel, 2018, Updegrove *et al*., 2016)). To determine whether Hfq had any impact on AzuC, AzuC-SPA levels were compared in wild type and Δ*hfq* cells grown to stationary phase under the same conditions as above (Fig 1C). AzuC-SPA levels were elevated in the Δ*hfq* strain for the cells grown in M63 galactose. This observation is consistent with potential posttranscriptional repression by Hfq and base-pairing sRNAs. However, before delving further into the regulation of AzuC expression, we wanted to learn more about the function of this 28 amino acid protein.

### AzuC protein is localized to the membrane

Information about the subcellular localization of proteins can give clues about possible interacting partners and functions in the cell. Secondary structural predictions suggested that AzuC has the potential to fold into an amphipathic helix (Fig 2A), indicating the protein might associate with the membrane. To test this, AzuC-SPA cells grown in M63 glucose to OD_600_ ∼1.0 were lysed, and cell extracts were homogenized and fractionated into soluble, inner membrane and outer membrane fractions by sucrose cushion fractionation (Fontaine *et al*., 2011, Rhoads *et al*., 1984). Consistent with the secondary structure prediction, immunoblot analysis of the fractions showed that AzuC-SPA was enriched in the inner membrane fraction, while the OmpA control protein was enriched in the outer membrane fraction (Fig 2B). Similar fractionation of untagged AzuC, expressed from a plasmid and detected by *α*-AzuC antiserum, also showed enrichment in the membrane fraction (Fig EV2A).

**Figure 2.**
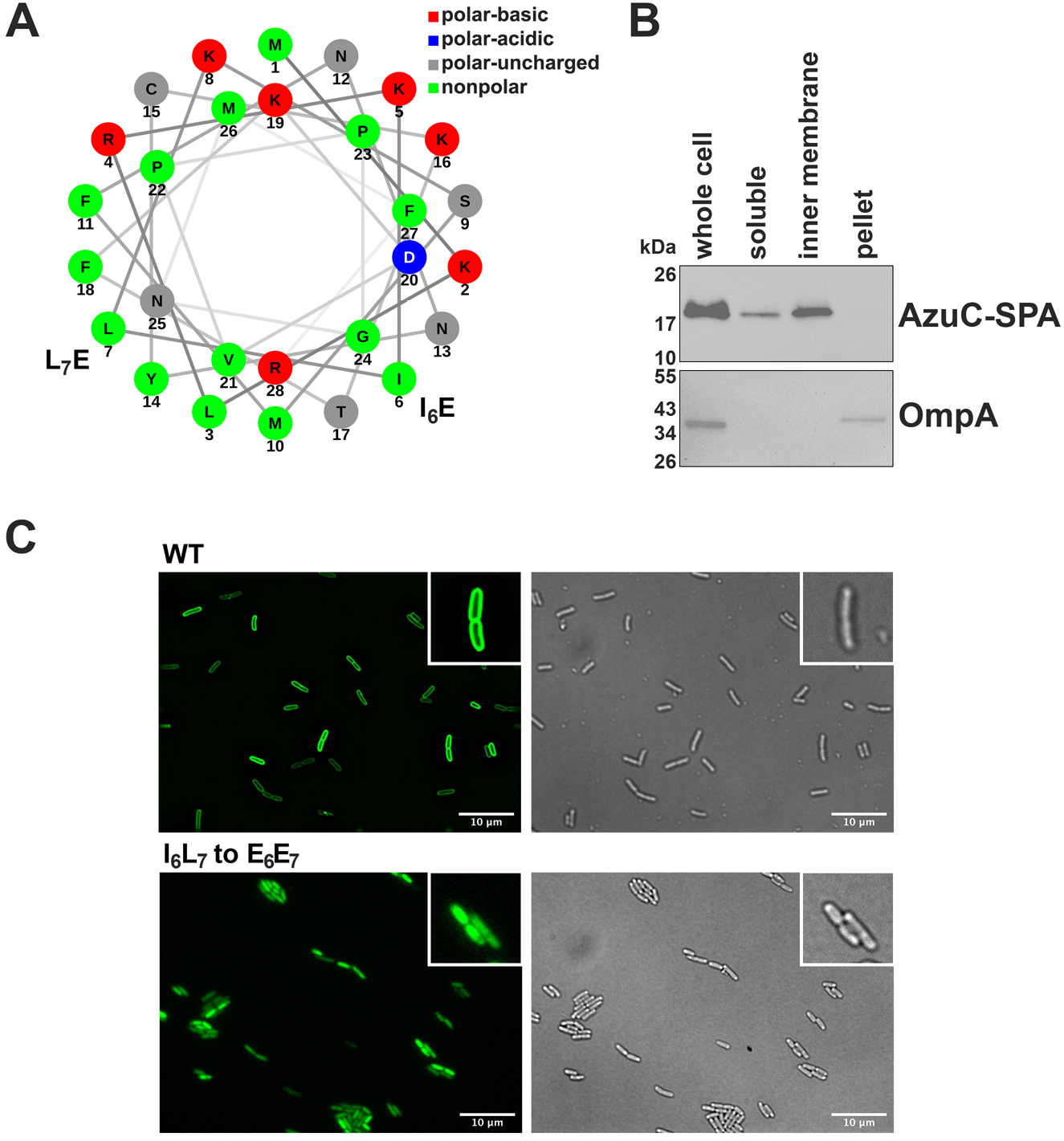
AzuC protein is membrane associated. A Helical wheel projection generated using NetWheels (Mól *et al*., 2019) showing amphipathic nature of AzuC. Mutations introduced in (C) are indicated. B Fractionation of AzuC-SPA strain. A culture expressing AzuC-SPA (GSO351) was grown in M63 glucose medium to OD_600_ ∼0.5, and cells were fractionated into a soluble, inner membrane, and pellet fractions, which were compared to the whole cell lysate. The top panel shows AzuC-SPA as detected with *α*-FLAG antibody. The bottom panel shows the outer membrane OmpA control detected with *α*-OmpA antibody. C Microscopy of AzuC-GFP. AzuC-GFP (GSO1008) and AzuC_I6L7 to E6E7_-GFP mutant (GSO1009) cells were grown in M63 glucose medium to OD_600_ ∼0.5 to observe membrane localization by fluorescent microscopy. Left panels are fluorescent images showing GFP labeled AzuC, and the right panels are the corresponding brightfield images. Insets provide an enlargement of a few cells.

The localization of AzuC to the membrane was further confirmed by fluorescence microscopy imaging of chromosomally-expressed AzuC C-terminally tagged with the green fluorescent protein (GFP) (Fig 2C). While wild type AzuC-GFP showed clear membrane localization, mutations replacing hydrophobic residues with charged residues (I_6_L_7_ to E_6_E_7_) eliminated the membrane localization. Taken together, these data support the hypothesis that AzuC is associated with the membrane as an amphipathic protein.

### AzuC protein interacts with GlpD

To further investigate the role of AzuC in the cell, we carried out co-purification assays to identify interacting proteins. Cells expressing chromosomally-encoded AzuC-SPA or previously-characterized AcrZ-SPA (Hobbs *et al*., 2012) as a control were grown in M63 glucose medium. Cell lysates prepared from exponentially-growing cells were applied to calmodulin beads, and the eluants from each column were separated by SDS-PAGE (Fig 3A). Unique bands from each of the elutions were sent for mass spectrometric analysis for identification. In two independent experiments, a prominent band of ∼60 kDa observed only for the AzuC-SPA cells was identified as the aerobic glycerol 3-phosphate dehydrogenase (GlpD), which catalyzes the oxidation of glycerol 3-phosphate. The most prominent band in the AcrZ sample was AcrB, a known interactor (Hobbs *et al*., 2012).

**Figure 3.**
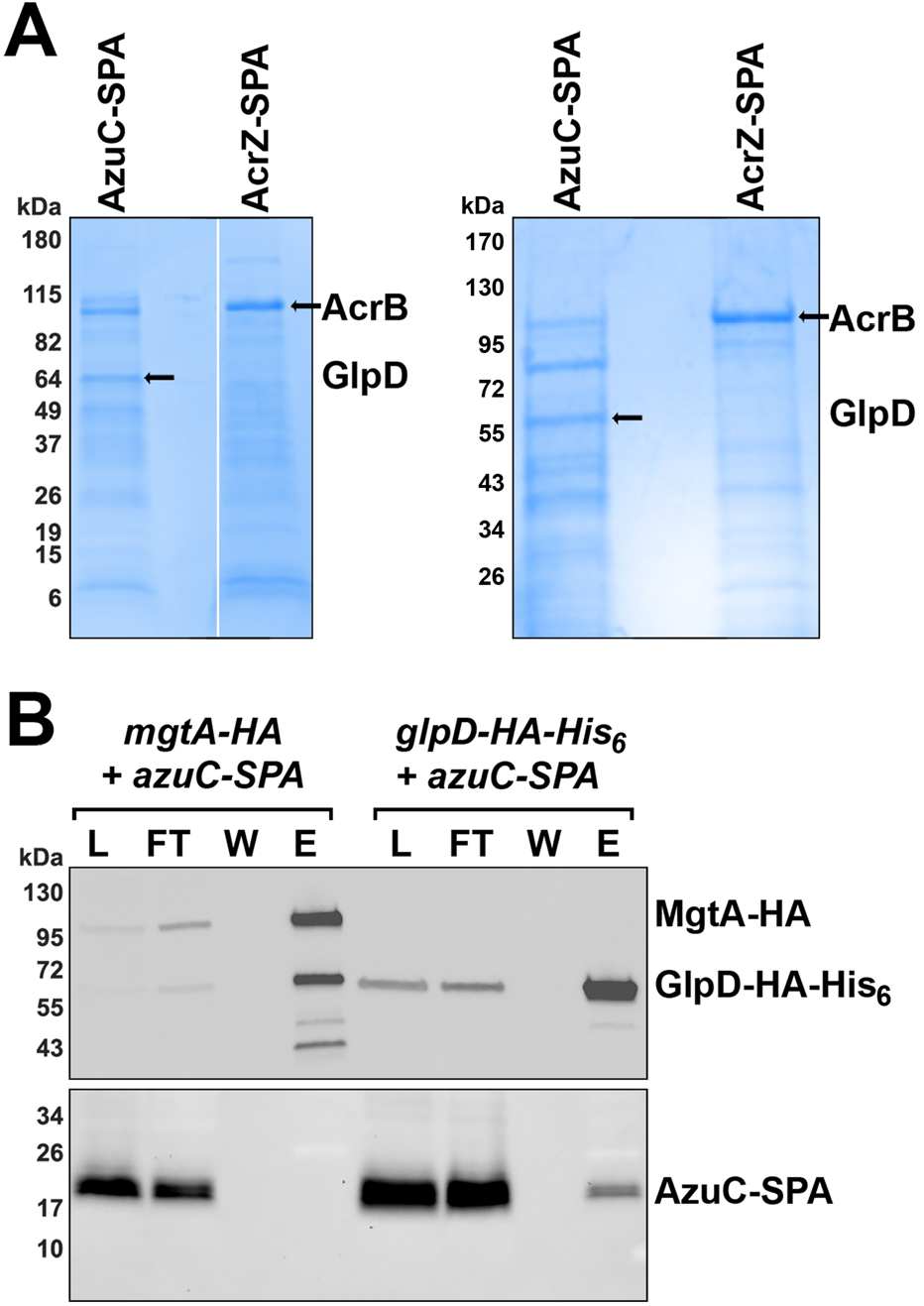
AzuC copurifies with GlpD. A GlpD co-purifies with AzuC-SPA. Cells expressing AzuC-SPA (GSO351) or AcrZ-SPA (GSO350) from the chromosome were grown in M63 glucose medium to OD_600_ ∼1.0 or in LB to OD_600_ ∼0.6, respectively. The cell lysates were split and passed over calmodulin beads. Eluants from each column were subjected to SDS-PAGE followed by Coomassie blue staining. The bands enriched in the eluant from the calmodulin beads and indicated by the arrows were excised from the gel and identified by mass spectrometry. B AzuC-SPA co-purifies with GlpD-HA-His_6_. Cells expressing either AzuC-SPA (GSO351) or GlpD-HA-His_6_ (GSO1011) from the chromosome were grown in M63 glucose or M63 glycerol media, respectively, to OD_600_ ∼1.0 and mixed in a 1:1 ratio. As a control, cells expressing MgtA-HA (GSO785) grown in N medium supplemented without added MgSO_4_ to OD_600_ ∼0.5, were mixed with the AzuC-SPA (GSO351) cells in the same ratio. The mixed cells were homogenized, cell lysates (L) were applied to *α*-HA beads and the flow-through (FT) samples were collected. The beads were washed (W), after which the bound proteins were eluted (E) and examined on immunoblots using either *α*-HA antibodies to detect MgtA-HA or GlpD-HA-His_6_ (top panel) or *α*-FLAG antibodies to detect AzuC (bottom panel).

We tested the interaction between AzuC and GlpD, by assessing reciprocal co-purification of AzuC-SPA with GlpD-HA-His_6_. Cells with chromosomally-encoded AzuC-SPA, grown to exponential phase in M63 glucose medium, were mixed with cells with chromosomally-encoded GlpD-HA-His_6_ grown to exponential phase in M63 glycerol medium, a condition where GlpD is known to be expressed. The mixed cells were lysed and incubated with dodecyl β-D-maltoside (DDM) to facilitate mixing of the membrane fractions. The mixed lysate was then applied to *α*-HA beads, washed and eluted (Fig 3B). As controls, similar purifications were carried out by mixing the AzuC-SPA cells with cells lacking tagged proteins grown in M63 glycerol medium (Fig EV2B) or cells expressing chromosomally-encoded MgtA-HA grown in N medium without added MgSO_4_ to induce MgtA expression (Fig 3B). The eluates were analyzed for the respective tagged proteins by immunoblot analysis by using *α*-FLAG and *α*-HA antibodies. Consistent with the first purification, AzuC-SPA co-purified with GlpD-HA-His_6_ and not with MgtA-HA, supporting the conclusion that GlpD is an interacting partner of AzuC. As we suggest for AzuC, GlpD has been reported to be a peripheral membrane protein that associates with the membrane through an amphipathic helix (Walz *et al*., 2002).

### AzuC protein increases GlpD activity

Binding of AzuC to GlpD could potentially impact the stability, localization, or activity of the enzyme as has been found for other small proteins (Hobbs *et al*., 2012, Wang *et al*., 2017). To distinguish among these possibilities, we first examined the levels of chromosomally-encoded GlpD-HA-His_6_ in cells transformed with pKK, pKK-AzuC, or pKK-AzuC_L3STOP_. In the latter two plasmids, the wild type or mutant (harboring a stop codon mutation of the third codon) *azuC* ORF was cloned downstream of the heterologous P_tac_ promoter and ribosome binding site on the pKK177-3 (pKK) plasmid. Cells were grown in M63 glucose medium to OD_600_ ∼1.0 and then transitioned to glycerol (pH 5.5) for 3 h given that chromosomally-expressed AzuC-SPA levels are elevated under these conditions (Fig 1B). The GlpD-HA-His_6_ protein levels were similar for all three strains grown under these conditions (Fig EV3A).

To test whether AzuC affects GlpD activity, we employed a dehydrogenase activity assay in which glycerol-3-phosphate oxidation to dihydroxyacetone phosphate (DHAP) is coupled to the reduction of yellow 2-(4,5-dimethyl-2-thiazolyl)-3,5-diphenyl-2*H*-tetrazolium bromide (MTT) to blue formazan, which is detected at OD_570_ (Wegener *et al*., 2016). Dehydrogenase activity based on this assay was found to be almost 2-fold lower in the absence of AzuC when extracts were made from a WT or a Δ*azuC* strain grown in M63 glucose medium and shifted to glycerol (pH 5.5) for 3 h. In contrast, overexpression of wild type AzuC, but not AzuC_L3STOP_, led to an increase in the dehydrogenase activity for the extracts (Fig 4A, middle panel). To further verify that this increase was GlpD dependent, the assay was repeated in a *ΔazuC ΔglpD* double mutant. The double mutant did not show the increase in dehydrogenase activity upon AzuC overexpression (Fig 4A, bottom panel), indicating that the interaction of the small protein AzuC with GlpD increases the dehydrogenase activity of the larger protein. Interestingly, we observed a GlpD-dependent decrease in dehydrogenase activity with the pKK-AzuC_L3STOP_ plasmid. We think this may be due to regulatory activity of the RNA (see below). The dehydrogenase assay also was carried out with these strains shifted to M63 glycerol (pH 7.0) where we observed similar, albeit somewhat smaller, effects of Δ*azuC* and AzuC overexpression on activity (Fig EV3B).

**Figure 4.**
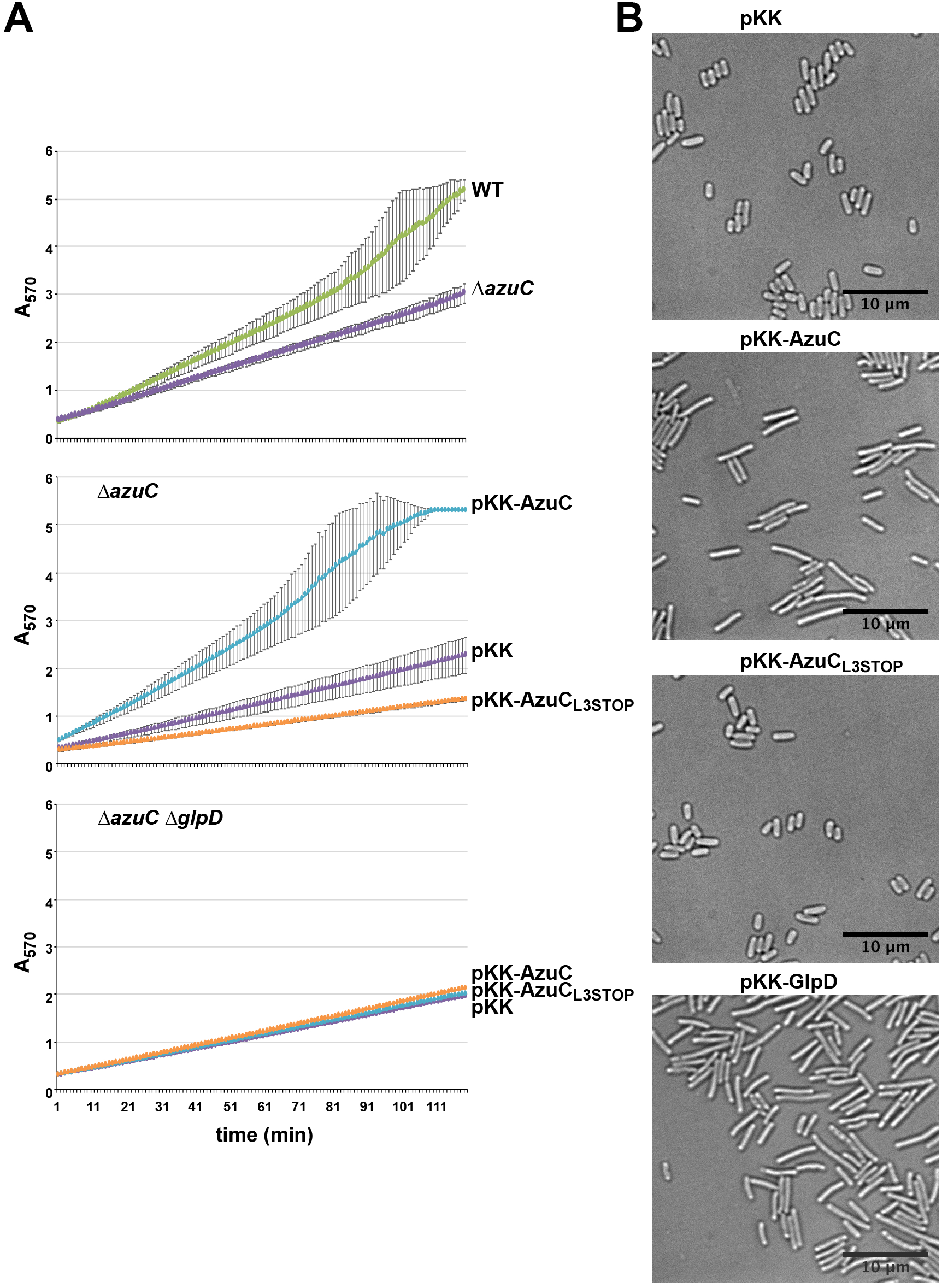
AzuC increases GlpD activity and affects cell shape. A Effect of AzuC overexpression on GlpD activity. WT or *ΔazuC::kan* cells (top panel), *ΔazuC::kan* (GSO193) (middle panel) or *ΔazuC ΔglpD::kan* (GSO1015) (bottom panel) cells transformed with pKK, pKK-AzuC, and pKK-AzuC_L3STOP_ were grown in M63 glucose medium to OD_600_ ∼1.0. Cells were washed and resuspended in M63 glycerol medium, pH 5.5 for 3 h prior to incubation with MTT and measurement of A_570_ reflecting the reduction of MTT to formazan, which is coupled to the oxidation of glycerol-3-phosphate to DHAP. B Effect of AzuC overexpression on *E. coli* cell morphology. *ΔazuC::kan* (GSO193) transformed with pKK, pKK-AzuC, pKK-AzuC_L3STOP_, or pKK-GlpD were grown in M63 glucose medium to OD_600_ ∼1.0. Cells were washed and resuspended in M63 glycerol medium, pH 5.5 for 3 h prior to microscopy.

### AzuC overexpression causes an increase in cell length

The substrate for GlpD, glycerol-3-phosphate, is a precursor for phospholipid biosynthesis. Thus, we wondered whether increasing the activity of GlpD by AzuC might bias the flow of glycerol-3-phosphate towards glycerol metabolism rather than phospholipid biosynthesis, which could impact cell morphology. To assess this, we carried out live-cell phase contrast microscopy of cells carrying pKK, pKK-AzuC or pKK-AzuC_L3STOP_ (Fig 4B). We observed AzuC overexpressing cells, but not those carrying the vector or pKK-AzuC_L3STOP_, had an elongated morphology. The elongated morphology was similar to the morphology observed for cells upon GlpD overexpression (Fig 4B) as well as the morphology reported for cells lacking phosphatidylethanolamine (Rowlett *et al*., 2017), which comprises ∼75% of the membrane phospholipid.

### AzuC and GlpD protein levels are repressed by the FnrS small RNA

We previously found that AzuC levels are higher under aerobic compared to anaerobic conditions (Hemm *et al*., 2010). Similarly, GlpD is required under aerobic conditions and is down-regulated during anaerobic growth while a second glycerol dehydrogenase, GlpABC, is required under anaerobic conditions. It was interesting to note that interactions between the anaerobic-induced sRNA FnrS and both the *azuC* and *glpD* mRNAs were found in genome-wide assays of RNA-RNA interactions on the Hfq chaperone (Melamed *et al*., 2020, Melamed *et al*., 2016). We also could predict base pairing between the 5’ end of FnrS and *azuC* as well as *glpD* (Fig 5A and 5C). These observations suggested possible FnrS-mediated repression of AzuC and GlpD synthesis. Consistent with this hypothesis, we observed higher AzuC-SPA levels in a Δ*fnrS* strain (Fig EV4A) and lower AzuC-SPA levels upon overexpression of WT FnrS and previously-generated FnrS-I and FnrS-II mutants (Durand & Storz, 2010) but not FnrS-III for which base pairing is predicted to be disrupted (Fig 5B). We similarly observed that WT FnrS, FnrS-I and FnrS-II, but not FnrS-III repressed a *azuC-lacZ* translational fusion expressed from the heterologous P_BAD_ promoter (Fig EV4B). Repression was restored for the FnrS-III mutant but not WT FnrS, FnrS-I, and FnrS-II by compensatory mutations in the *azuC-lacZ*-III mutant fusion demonstrating direct base-pairing between FnrS and the *azuC* mRNA. We also observed slightly lower levels of the GlpD-HA-His_6_ protein upon overexpression of WT FnrS, FnrS-I and FnrS-II, but not FnrS-III (Fig 5D). Together these results indicate that the 5’ end of FnrS base pairs with the *azuC* and *glpD* mRNAs to repress synthesis of the two proteins.

**Figure 5.**
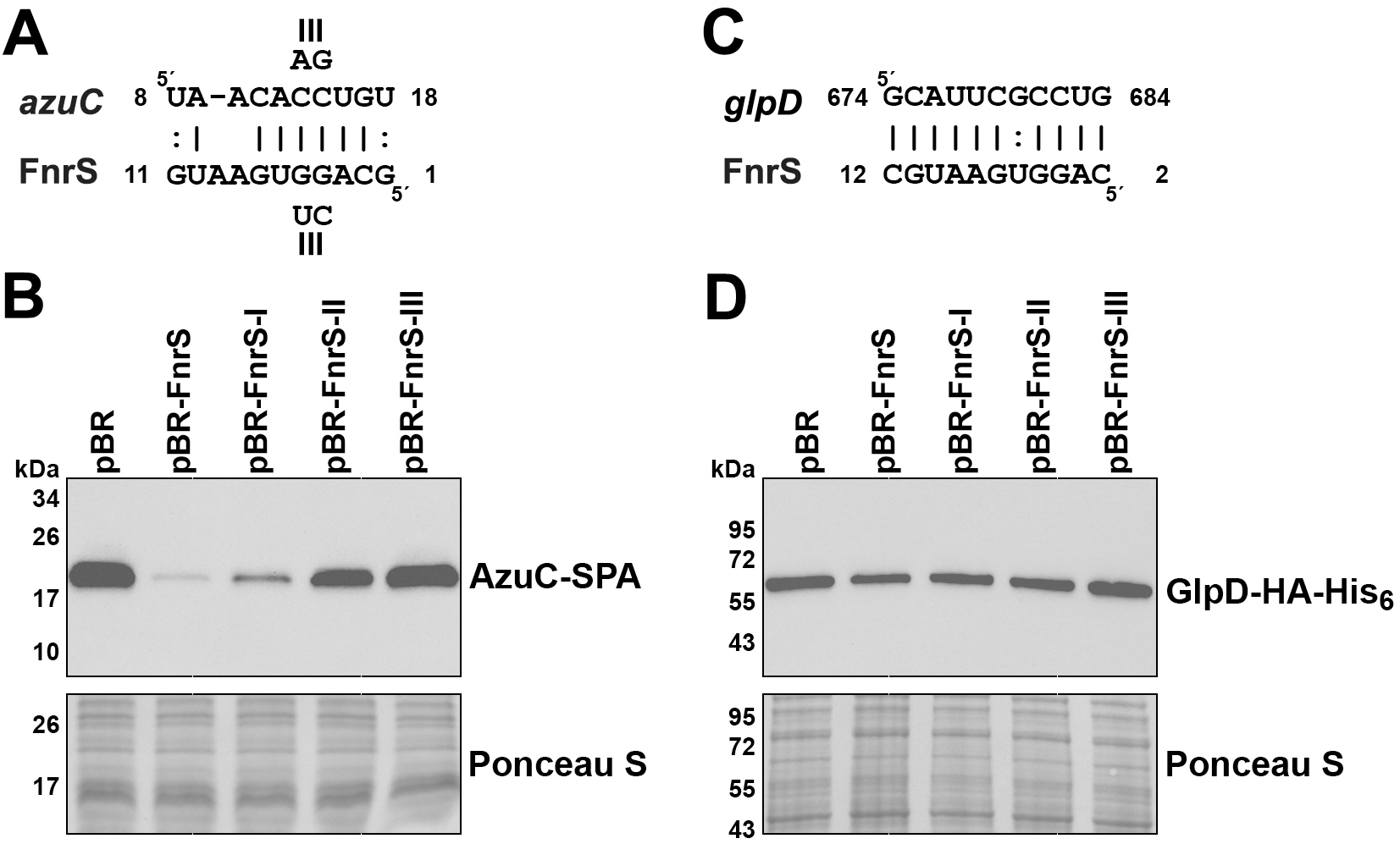
FnrS sRNA represses synthesis of both AzuC and GlpD. A Predicted base pairing between FnrS and *azuC*. The coordinates for both are relative to the +1 of the transcript. B Effect of FnrS overexpression on AzuC-SPA levels. Cultures of the *azuC-SPA::kan* (GSO351) strain carrying pBR, pBR-FnrS, pBR-FnrS-I, pBR-FnrS-II, or pBR-FnrS-III were grown in LB with 1 mM IPTG to OD_600_ ∼0.5. *α*-FLAG antibodies were used to detect the SPA tag. C Predicted base pairing between FnrS and *glpD*. The predicted region of pairing in *glpD* is within the coding sequence. The coordinates for FnrS are relative to the +1 of the transcript, while the coordinates for *glpD* are relative to the first nucleotide of the start codon. D Effect of FnrS overexpression on GlpD-HA-His_6_ levels. Cultures of the *glpD-HA-His_6_* (GSO1011) strain carrying pBR, pBR-FnrS, pBR-FnrS-I, pBR-FnrS-II, or pBR-FnrS-III were grown in LB with 1 mM IPTG to OD_600_ ∼0.5. *α*-His antibodies were used to detect GlpD-HA-His_6_. For (B) and (D), the membrane was stained with Ponceau S stain to control for loading.

### AzuCR overexpression reduces growth in glycerol and galactose

Given the AzuC effect on GlpD together with the different AzuC levels for cells grown in the presence of different carbon sources, we examined the consequences of AzuC overexpression from the pKK vector for growth in glucose and glycerol at pH 7.0 and 5.5 and galactose at pH 7.0 (Fig 6 and Fig EV5A). For pKK-AzuC cells, but not the pKK vector control and pKK-AzuC_L3STOP_ cells, we observed a significant growth defect in M63 glycerol pH 5.5 and a partial defect in M63 glycerol pH 7.0, consistent with the larger effect of AzuC on GlpD activity in M63 glycerol pH 5.5 compared to pH 7.0. A similar phenotype was observed for overexpression of AzuC-SPA indicating that the tagged derivative of AzuC is functional (Fig EV5B). Growth in minimal medium with either glucose or galactose was not significantly changed by the pKK-AzuC plasmid.

**Figure 6.**
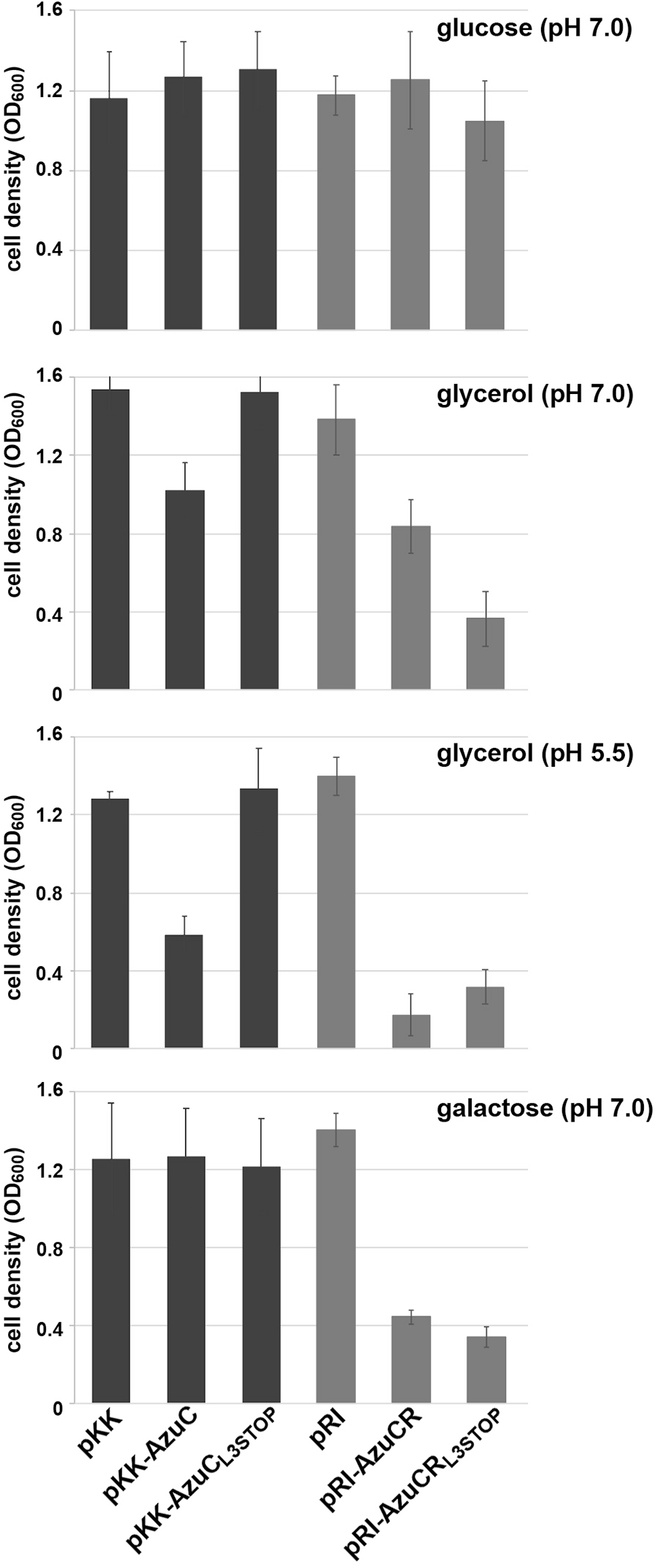
AzuC and AzuR overexpression leads to different growth phenotypes in different carbon sources. Growth of the *ΔazuC::kan* strain (GSO193) transformed with pKK, pKK-AzuC, pKK-AzuC_L3STOP_, pRI, pRI-AzuCR, or pRI-AzuCR_L3STOP_ in M63 medium with different carbon sources, glucose (pH 7.0), glycerol (pH 7.0 and 5.5), and galactose (pH 7.0), was measured 16 h after dilution by OD_600_. The full growth curves are in Fig EV5A.

We also examined the effect of overexpressing the full-length *azuC* mRNA (pRI-AzuCR) without or with the L3STOP mutation (pRI-AzuCR_L3STOP_) (Fig 6). Interestingly, we observed different effects on growth for these plasmids. While growth in M63 glucose pH 7.0 was not affected, pRI-AzuCR led to a growth defect in M63 glycerol pH 7.0 and even more so at pH 5.5 as well as in M63 galactose pH 7.0. Contrary to the detrimental effect of the L3STOP mutation when only the *azuC* coding sequence was included, the L3STOP mutation in the full-length transcript still blocked growth and, in M63 glycerol pH 7.0, actually exacerbated the growth defect. This observation suggested the transcript could have a second role as a regulatory RNA, which we have denoted AzuR.

### AzuR functions as an sRNA to repress *cadA* and *galE*

Based on our findings that the AzuCR transcript could have a second role as an sRNA, we investigated its potential as a base-pairing sRNA by searching for possible base pairing targets using TargetRNA2 (Kery *et al*., 2014) and IntaRNA (Mann *et al*., 2017) prediction programs. Given the reduced growth associated with AzuCR overexpression in cells grown in galactose and low pH, we focused on potential target genes that might be important under these conditions. One predicted target with extensive potential base pairing was *cadA*, encoding lysine decarboxylase (Fig 7A). Consistent with AzuR-mediated regulation of *cadA*, we observed decreased expression of a *cadA-gfp* fusion upon AzuCR overexpression and even more so for AzuCR_L3STOP_ overexpression (Fig 7B). Additionally, there were higher overall levels of *cadA-gfp* expression in the Δ*azuC* strain compared to the WTstrain, suggesting that chromosomally-encoded AzuC contributes to the repression. Consistent with the base pairing predicted in Fig 7A, the M1 mutations in AzuCR_L3STOP_ reduced *cadA-gfp* repression, while regulation was restored when compensatory mutations were introduced in the *cadA-gfp* construct (Fig 7C). Another predicted target for base pairing with AzuR was *galE* (Fig 7D), the first gene in the *galETKM* galactose operon. The AzuCR_L3STOP_ derivative also repressed a *galE-gfp* fusion in both the WT and Δ*azuC* backgrounds, with partial repression by AzuCR (Fig 7E). Again there is direct base pairing between AzuCR and *galE*, as the M2 mutations in AzuCR_L3STOP_ or *galE* alone reduced AzuCR-mediated *galE-gfp* repression, while repression was restored when both compensatory mutations were present (Fig 7F).

**Figure 7.**
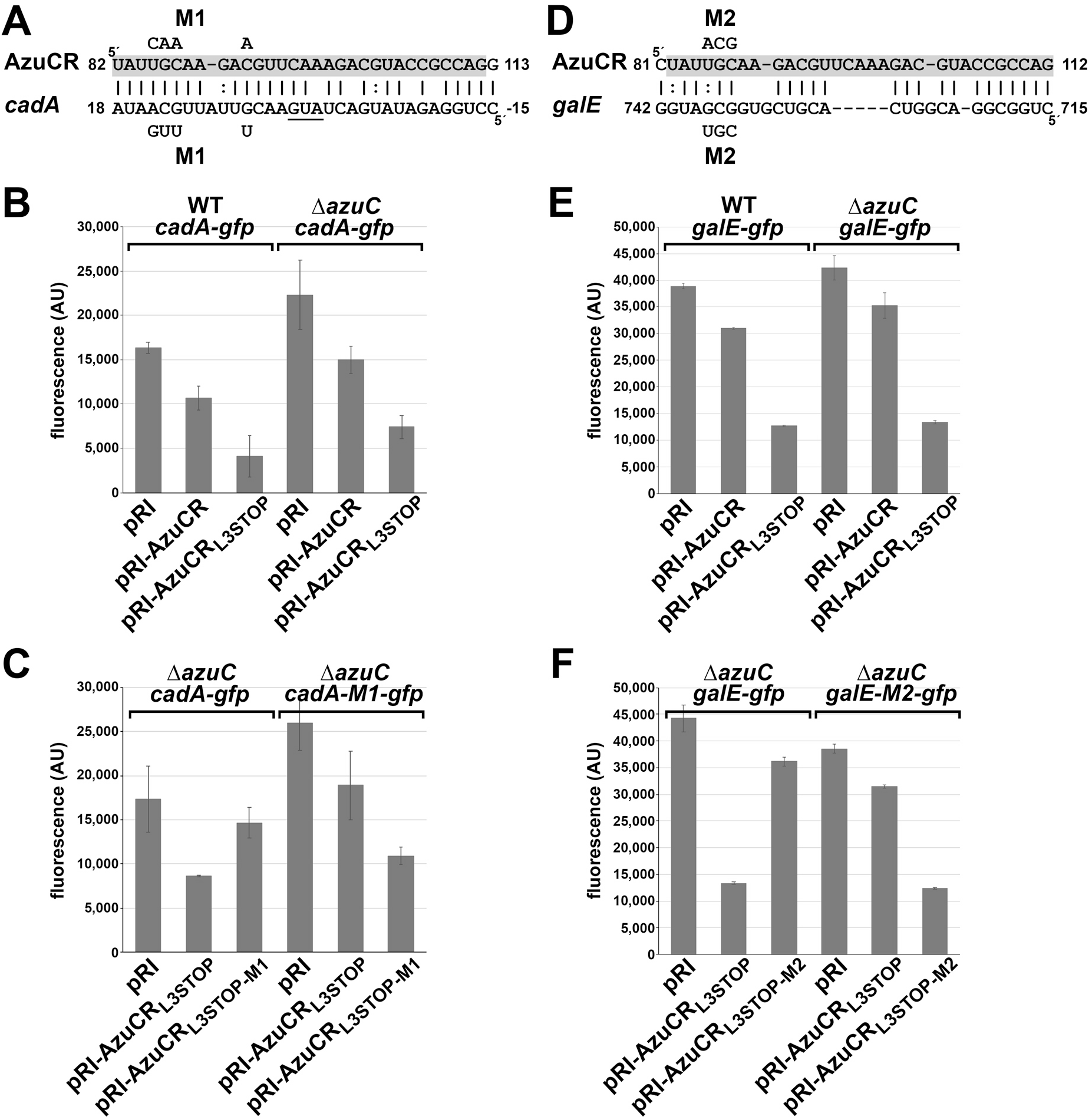
AzuR represses *cadA* and *galE* expression. A AzuCR-*cadA* base pairing predicted by TargetRNA2 (Kery *et al*., 2014). The coordinates for AzuCR are relative to the +1 of the transcript, while the coordinates for *cadA* are relative to the first nucleotide of the start codon. Mutations introduced into AzuCR and *cadA* are indicated. B Effect of AzuCR and AzuCR_L3STOP_ overexpression on a *cadA-gfp* fusion in wild type and Δ*azuC* backgrounds. WT and Δ*azuC::kan* (GSO193) cells were co-transformed with a reporter plasmid expressing a *cadA-gfp* translational fusion and either the empty pRI vector, AzuCR, or AzuCR_L3STOP_. C Test of AzuCR-*cadA* base pairing. Δ*azuC::kan* (GSO193) cells were cotransformed with the WT *cadA-gfp* translational fusion reporter plasmid or a M1 derivative with mutations in the predicted region of base pairing along with the empty pRI vector, AzuCR_L3STOP_, or mutant AzuCR_L3STOP-M1_. The mutations in the *cadA-gfp* translational fusion and AzuCR_L3STOP_ are indicated (A). D AzuCR-*galE* base pairing predicted by IntaRNA (Mann *et al*., 2017). The coordinates for AzuCR are relative to the +1 of the transcript, while the coordinates for *galE* are relative to the first nucleotide of the start codon. Mutations introduced into AzuCR and *galE* are indicated. E Effect of AzuCR and AzuCR_L3STOP_ overexpression on a *galE-gfp* fusion in wild type and Δ*azuC* backgrounds. WT and Δ*azuC::kan* (GSO193) cells were co-transformed with a reporter plasmid expressing a *galE-gfp* translational fusion and either the empty pRI vector, AzuCR or AzuCR_L3STOP_. F Test of AzuCR-*galE* base pairing. Δ*azuC::kan* (GSO193) cells were cotransformed with the WT *galE-gfp* translational fusion reporter plasmid or a M2 derivative with mutations in the predicted region of base pairing along with the empty pRI vector, AzuCR_L3STOP_, or mutant AzuCR_L3STOP-M2_. The mutations in the *galE-gfp* translational fusion and AzuCR_L3STOP_ are indicated in (D). For (B), (C), (E) and (F), cells were grown in LB for 3 h before measuring the fluorescence corresponding to GFP expression. The average of three independent trials is shown, and the error bars represent one SD.

### AzuCR RNA association with Hfq and ProQ is not required for *cadA* repression

Consistent with the observation that Δ*hfq* impacts AzuC protein levels (Fig 1B), we found that the AzuCR mRNA co-immunoprecipitates with Hfq (Fig 8A). Another RNA chaperone that has been found to bind to sRNA-mRNA pairs in *E. coli* and impacts the stability of some RNAs is ProQ (Melamed *et al*., 2020). As with Hfq, AzuCR co-immunoprecipitates with ProQ (Fig 8A). However, in contrast to the increased AzuCR RNA levels in the Δ*hfq* background, AzuCR RNA levels were decreased in the Δ*proQ* background. We wondered whether AzuCR functions as an sRNA repressor were mediated by Hfq or ProQ and examined repression of the *cadA-gfp* in Δ*hfq* and Δ*proQ* single as well as Δ*hfq* Δ*proQ* double mutant backgrounds. GFP activity levels overall were lower when Hfq was absent, but we observed *cadA-gfp* repression by AzuCR_L3STOP_ overexpression in all backgrounds (Fig 8B). These observations indicate that although both Hfq and ProQ bind to AzuCR, the RNA chaperones are not required for the repression of *cadA* when AzuCR_L3STOP_ is overexpressed, possibly due to the long region of potential base pairing.

**Figure 8.**
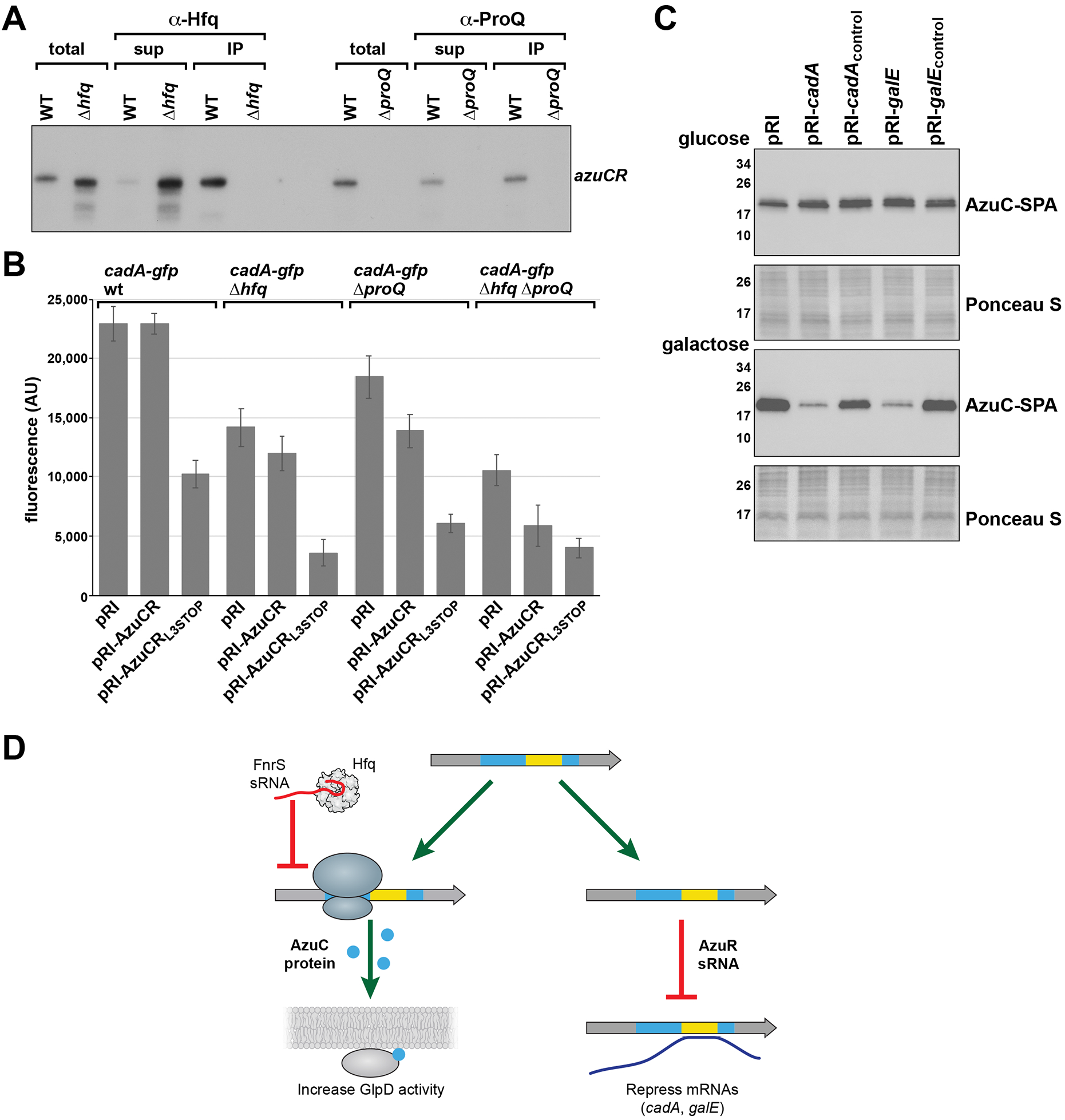
AzuCR mRNA and base pairing activities are differentially affected by Hfq and ProQ. A AzuCR co-immunoprecipitation with Hfq and ProQ. Extracts from MG1655 (GSO982), Δ*hfq-cat::sacB* (GSO954) and Δ*proQ::kan* (GSO956) cells grown in M63 glucose were incubated with *α*-Hfq or *α*-ProQ antiserum. Total and RNA chaperone-bound RNA was extracted and subjected to Northern analysis using an oligonucleotide probe specific for AzuCR. B Effect of Δ*hfq::kan* (GSO955), Δ*proQ::kan* (GSO956) and Δ*hfq* Δ*proQ::kan* (GSO959) double mutant on AzuCR repression of *cadA-gfp*. *cadA-gfp* expression from pXG10-SF in the presence of AzuCR or AzuCR_L3STOP_ in WT, Δ*hfq* or Δ*proQ* backgrounds. The average of three independent trials is shown, and the error bars represent one SD. C Effect of *cadA*_base pairing_, *cadA*_control_, *galE*_base pairing_ and *galE*_control_ on AzuC-SPA levels in cells (GSO351) transformed with the respective overexpression plasmid and grown in M63 medium supplemented with glucose or galactose. Samples were taken at OD_600_ ∼0.5 and *α*-FLAG antibody was used to detect the SPA tag. The membranes stained with Ponceau S stain serves as a loading control. D Model for the different functions of the AzuCR RNA. For growth in M63 glycerol, pH 5.5, the RNA can be translated to give the 28 amino acid amphipathic AzuC protein, which increases the activity of GlpD glycerol-3-phosphate dehydrogenase. Under anaerobic conditions this translation and the translation of GlpD is blocked by the FnrS sRNA. The RNA can also act as the AzuR base-pairing sRNA to repress synthesis of CadA and GalE.

### AzuC translation and AzuR base pairing activity interfere

The region of base pairing between AzuR and *cadA* and *galE* (89-107 nt relative to the transcription start) overlaps the *azuC* coding sequence (40-126 nt relative to the transcription start) raising the question of whether the mRNA and base pairing activities of the AzuCR RNA interfere which each other. The hypothesis that translation interferes with base pairing is supported by the observations that AzuCR_L3STOP_ was more effective at repressing the *cadA-gfp* and *galE-gfp* fusions than AzuCR (Fig 7B and 7E). To determine if base pairing activity reciprocally interfers with translation, we examined the levels of chromosomally-encoded AzuC-SPA upon overexpression of the base pairing regions of *cadA* and *galE* along with control regions of these genes not predicted to base pair with AzuCR. Interestingly, no repression was observed for cells grown in M63 glucose. In contrast, the base pairing fragments, but not the control fragments, led to decreased AzuC-SPA levels for cells grown in M63 with galactose (Fig 8C). These observations suggest that base pairing can interfere with translation, particularly when AzuC protein levels overall are low as is the case for cells grown in M63 galactose.

## Discussion

Successful adaptation to varying environmental conditions requires regulation that can rapidly lead to changes in metabolism. Along with transcription factors, sRNAs and small proteins are emerging as important regulators. While base-pairing sRNAs generally are not thought to be translated, a few have been reported to encode small proteins. Even fewer of these dual-function RNAs have been characterized. Here we report that the 164 nt RNA previously reported to encode a 28 amino acid small protein (AzuC) (Hemm *et al*., 2010) also functions as a regulatory RNA (AzuR). We show the AzuC protein binds and activates GlpD, an enzyme at the junction of respiration, glycolysis, and phospholipid biosynthesis (Fig 3A and 3B), while the RNA base pairs with and represses expression from the *cadA* and *galETKM* mRNAs (Fig 7).

### AzuCR is a unique dual-function RNA

There are a number of features of AzuCR that are unique compared to other dual-function RNAs. First, while the regions involved in base pairing and protein coding are separate for the well-characterized *E. coli* SgrS-SgrT and *Staphylococcus aureus* RNAIII RNAs (reviewed in (Raina *et al*., 2018)), as well as the *Vibrio cholerae* VcdRP RNA described in a co-submitted paper (Venkat *et al*., 2021), the region of AzuCR involved in base pairing with the *cadA* and *galE* targets overlaps the *azuC* coding sequence (Fig 1A). Another notable feature of this dual-function RNA is that while the gene is not broadly conserved, the levels of the RNA and protein are highly regulated. In our previous study, we observed the AzuC protein accumulates in minimal glucose medium as well as in response to low pH, high temperature, and hydrogen peroxide, while the levels are low under anaerobic conditions (Hemm *et al*., 2010). The regulation in response to glucose availability was shown to be at the transcriptional level via CRP derepression. In our current study, we show that AzuC repression under anaerobic conditions is mediated by the Hfq-dependent sRNA FnrS, which base pairs near the AzuC ribosome binding site (Fig 5A). To the best of our knowledge, FnrS regulation of AzuCR is the first example of another sRNA regulating the translation of a dual-function RNA. The observation that AzuC levels are higher in Δ*hfq* compared to Δ*fnrS* mutant cells (Fig EV4A) suggests that other Hfq-dependent sRNAs might repress AzuC synthesis. This is further indicated by the discordance between RNA and protein levels under some conditions (Fig 1B).

Consistent with the discordant expression of the AzuCR RNA compared to the AzuC protein, we found that the small protein and base pairing activities modulate overlapping but distinct pathways; AzuC plays a role in glycerol metabolism (Fig 3 and 4) and AzuCR impacts galactose and glycerol metabolism (Fig 6). The regulation of different pathways by the two activities of AzuCR contrasts with SgrST RNA where the SgrT protein and the base-pairing SgrS RNA both down-regulate the PtsG glucose transporter activity.

### AzuC stimulation of GlpD activity

Although the functions of only a few small proteins have been described, most are inhibitory (reviewed in (Storz *et al*., 2014)). Thus, AzuC is unusual in that it increases GlpD activity. GlpD, one of the key flavin-linked primary dehydrogenases of the respiratory electron transport chain, catalyzes the oxidation of glycerol-3-phosphate to DHAP (Yeh *et al*., 2008). GlpD exists in in both soluble and membrane-bound forms and is only fully active when the enzyme is associated with the cytoplasmic membrane through lipid-enzyme interactions or when reconstituted with phospholipids *in vitro* (Yeh *et al*., 2008, Schryvers *et al*., 1978, Robinson & Weiner, 1980). Interestingly, GlpD activity was previously reported to be increased by amphipaths (Robinson & Weiner, 1980). Like GlpD, AzuC is an amphipathic protein localized to the cytoplasmic membrane (Fig 2). Thus, it is possible that AzuC promotes GlpD binding to the cytoplasmic membrane. AzuC also could change the stability of GlpD, though we did not observe obvious differences in protein levels (Fig EV3A). Alternatively, AzuC could increase GlpD activity by causing a conformational change. Interestingly, the 29 amino acid *V. cholerae* VcdP similarly was found to increase citrate synthase activity, likely by counteracting the inhibitory effects of NADH (Venkat *et al*., 2021).

The physiological role of AzuC activation of GlpD, particularly at pH 5.5, is an interesting question. We suggest AzuC binds GlpD under acidic conditions to modulate the levels of glycerol-3-phosphate, which can undergo two acylation steps to form phosphatidic acid, a precursor for the phospholipids phosphatidylethanolamine (PE), phosphatidylglycerol (PG) and cardiolipin (CL). Bacterial adaptation to environmental stress can be accompanied by changes in the lipopolysaccharide (LPS) structure, the phospholipid composition, and the protein content of the inner and outer membranes (Rowlett *et al*., 2017). These changes in turn impact cell division, energy metabolism, osmoregulation as well as resistance to cationic antimicrobial peptides (CAMPs). In support of the hypothesis that AzuC activation of GlpD affects membrane composition, we observed that cells overexpressing AzuC or GlpD grown in low pH showed increased cell length (Fig 4B) and reduced growth (Fig 6), phenotypes that have also been observed in cells lacking the phospholipids PE and PG/CL (Rowlett *et al*., 2017).

### AzuR repression of the *cadA* and the *galETKM* mRNAs

We found that as a base pairing RNA, AzuR represses expression of CadA (Fig 7A, B, and C), which is induced under acidic growth conditions and confers resistance to weak organic acids produced during carbohydrate fermentation under anaerobiosis and phosphate starvation (reviewed in (Kanjee & Houry, 2013)). This regulation could partially explain the growth defect for cells growing in glycerol pH 5.5 observed upon AzuCR overexpression with and without a stop codon (Fig 6). We also identified *galETKM* mRNA as another direct AzuR target (Fig 7D, E, and F). Consistent with this regulation, we see a drastic growth defect with galactose as the sole carbon source upon overexpression of AzuCR with and without a stop codon (Fig 6). While AzuR base pairs near the ribosome binding site of the *cadA* mRNA likely blocking ribosome binding, the base pairing with the *galETKM* mRNA is internal to the *galE* coding sequence. We suggest that for this mRNA, base pairing may lead to changes in mRNA stability or alternatively the Rho-dependent transcription termination reported for the *galETKM* mRNA (Wang *et al*., 2014). Since we also observe an RNA-dependent growth phenotype for cells grown in glycerol pH 7.0, we suggest that AzuR might target other genes, particularly genes related to glycerol metabolism.

### Competition between two AzuCR activities

Several of our experiments indicate that there is competition between the mRNA and base pairing activities of AzuCR. We observed that a stop codon blocking translation improves AzuR base pairing activity (Fig 7B and 7E) and overexpression of fragments of the base pairing targets *cadA* and *galE* inhibits AzuC translation (Fig 8C). The conflict between base pairing and translation raises intriguing questions about what activity predominates under different growth conditions, whether the RNA can transition from one function to the other, what factors determine which activity predominates, and how the two activites evolved. We suggest that there are a number of scenarios for how AzuCR could act. There may be conditions where AzuCR acts solely a riboregulator and other conditions where AzuCR is solely an mRNA. There also may be conditions where there are two populations of AzuCR, some transcripts acting as an sRNA and others being translated. Additionally, AzuCR could first act as an mRNA and subsequently go on to act as a riboregulator.

The factors that regulate the distribution of AzuCR between these regulatory roles are not fully understood but clearly depend on the levels of the AzuCR RNA, sRNAs that repress AzuC translation, the levels of the mRNA targets of AzuCR, the levels of the Hfq and ProQ chaperones and likely other factors. The observed FnrS-dependent repression of AzuC synthesis is at least partially dependent on Hfq, while the stability of the RNA appears to depend on ProQ. The finding that AzuCR may only be a base-pairing RNA under specific conditions raises caveats for global approaches such as RIL-seq (Melamed *et al*., 2016) or RNA-seq after pulse overexpression for identifying sRNA targets and function. If these experiments are carried out under conditions where translation predominates, the effects of the base pairing activity may not be detected. These questions about AzuCR likely are relevant for other dual-function RNAs and are important directions for future research.

## Materials and Methods

### Bacterial strains and plasmid construction

Bacterial strains, plasmids, and oligonucleotides used in this study are listed in Appendix Tables S1, S2 and S3, respectively. *E. coli* strains are derivatives of wild-type MG1655 (F-lambda-*ilvG*-*rfb*-50 *rph*-1). Tagged strains were generated by λ Red–mediated recombineering (Yu *et al*., 2000) using NM400 and the oligonucleotides listed in Appendix Table S3. pJL148 (Zeghouf *et al*., 2004) was used as the template to amplify the SPA tag. The chromosomal P_BAD_-5’UTR*_azuC_*-*lacZ* and P_BAD_-5’UTR*_azuC_*-*lacZ* III fusions (carrying the first 87 nt of the *azuC* mRNA fused to the seventh codon of the *lacZ* coding sequence) were created by carrying out PCR using primers listed in Appendix Table S3 to applify the desired region of *azuC* followed by integration of the product into the chromosome of PM1205 (Mandin & Gottesman, 2009). Alleles marked by antibiotic markers were moved between strains by P1 transduction. When necessary, kanamycin resistance cassettes were excised from the chromosome by FLP-mediated recombination using the FLP recombinase encoded on pCP20 (Cherepanov & Wackernagel, 1995). All plasmids are derivatives of pAZ3 (Kawano *et al*., 2005), pKK177-3 (Brosius & Holy, 1984), pRI (Opdyke *et al*., 2004), pBRplac (Guillier & Gottesman, 2006) or pXG10-SF (Corcoran *et al*., 2012). All chromosomal mutations and fusions and plasmid inserts were confirmed by sequencing.

### Bacterial growth

Cells were grown in Luria-Bertani broth (LB) or M63 minimal media supplemented with 0.001% vitamin B1 and glucose, glycerol or galactose (0.2%, 0.4% or 0.2%, respectively). For some experiments, M63 medium was buffered to pH 5.5 with 100 mM MES. Cells were grown to the indicated OD_600_ after a 1:100 dilution of the overnight culture grown in LB, except for all M63 glycerol, pH 5.5 samples, where overnight cultures were grown in M63 glycerol, pH 5.5. Where indicated, media contained antibiotics with the following concentrations: ampicillin (100 μg/ml), chloramphenicol (25 µg/ml) and kanamycin (30 µg/ml).

### Immunoblot analysis

The cell pellet from 1 ml of cells grown in the indicated medium was resuspended in 1X PBS (KD Medical), 7 µl of 2X Laemmli buffer (BioRad) and 2 µl of β-mercaptoethanol, and 10 µl were loaded on a Mini-PROTEAN TGX 5%–20% Tris-Glycine gel (Bio-Rad) and run in 1X Tris Glycine-SDS (KD Medical) buffer. The proteins were electro-transferred to nitrocellulose membranes (Invitrogen) for 1 h at 100 V. Membranes were blocked with 5% non-fat milk (BioRad) in 1X PBS with 0.1% of Tween 20 (PBS-T) for 1 h and probed with a 1:3,000 dilution of *α*-FLAG-HRP antiserum (Sigma), 1:1,000 dilution of *α*-AzuC antiserum (New England Peptide); 1:1,000 dilution of *α*-His-HRP antiserum (Qiagen), or 1:1,000 dilution of *α*-OmpA antiserum (Antibody Research Corporation) in the same PBS-T buffer with 5% milk for 1 h. After the incubation with the *α*-AzuC and *α*-OmpA antiserum, membranes were incubated with a 1:2,000 dilution of HRP-labelled anti-rabbit antibody (Life Technologies). All blots were washed 4X with PBS-T and then developed with a Amersham ECL Western Blotting Detection Kit (GE Healthcare).

### Total RNA isolation

Cells corresponding to the equivalent of 10 OD_600_ were collected by centrifugation, and snap frozen in liquid nitrogen. RNA was extracted according to the standard TRIzol (Thermo Fisher Scientific) protocol. Briefly, 1 ml of room temperature TRIzol was add to cell pellets, resuspended thoroughly to homogenization, and incubated for 5 min at room temperature. After the addition of 200 µl of chloroform and thorough mixing by inversion, samples were incubated for 10 min at room temperature. After samples were centrifuged for 10 min at 4°C on maximal speed, the upper phase (∼0.6 ml) was transferred into a new tube and 500 µl of isopropanol was added. Samples again were mixed thoroughly by inversion, incubated for 10 min at room temperature and centrifuged at maximal speed for 15 min at 4°C. RNA pellets were washed twice with 75% ethanol and then dried at room temperature. RNA was resuspended in 20-50 µl of DEPC water and quantified using a NanoDrop (Thermo Fisher Scientific).

### Northern analysis

Total RNA (5-10 µg per lane) was separated on denaturing 8% polyacrylamide gels containing 6 M urea (1:4 mix of Ureagel Complete to Ureagel-8 (National Diagnostics) with 0.08% ammonium persulfate in 1X TBE buffer at 300 V for 90 min. The RNA was transferred to a Zeta-Probe GT membrane (Bio-Rad) at 20 V for 16 h in 0.5X TBE, UV-crosslinked, and probed with ^32^P-labeled oligonucleotides (Listed in Table 1) in ULTRAhyb-Oligo buffer (Ambion Inc.) at 45°C. Membranes were rinsed twice with 2X SSC-0.1% SDS at room temperature, once with 0.2X SSC-0.1% SDS at room temperature, washed for 25 min with 0.2X SSC-0.1% SDS at 45°C, followed by a final rinse with 0.2X SSC-0.1% SDS at room temperature before autoradiography was performed with HyBlot CL film (Denville Scientific Inc.).

### Sub-cellular fractionation

Cells with chromosomally-encoded AzuC-SPA were grown in the indicated medium at 37°C to an OD_600_ ∼0.3, centrifuged at 20,000 × g for 10 min at 4°C, resuspended in fractionation buffer (1/20 vol of 20% sucrose, 50 mM Tris pH 8) with 1 mM EDTA and 0.1 mg/ml lysozyme, and then incubated 1 h at 25°C with gentle shaking. After the cells were centrifuged at 20,000 × g for 15 min at 4°C, the top periplasmic fraction was removed. The pellet fraction was resuspended in water to lyse the spheroplasts. The resulting crude lysate was passed through a 30-gauge syringe needle 6X to homogenize the sample and reduce viscosity. The lysate was then clarified by centrifugation at 20,000 × g for 5 min at 4°C. This was repeated 3X. A 500 µl of the clarified lysate was layered on top of a 500 µl-sucrose cushion (5 mM EDTA and 1.4 M sucrose. Samples were centrifuged at 130,000 × g for 2 h at 4°C in a TLA100.3 rotor (Beckman Optima TLX table top centrifuge). Following centrifugation, 425 µl was carefully removed from the top layer (soluble fraction). Then, the interface and remaining liquid were removed (inner membrane fraction). The pelleted material was resuspended in 500 µl of fractionation buffer (pellet fraction). SDS was added to all fractions (final concentration 1%) and the samples were incubated overnight at room temperature. Equal volumes of fractions were assayed by immunoblotting with *α*-FLAG-HRP and *α*-OmpA antibody.

Cells expressing AzuC from a plasmid were grown as above, collected by centrifugation at 4K rpm for 10 min at 4°C, resuspended as above but incubated 10 min on ice. After the lysate was incubated as above the periplasmic fraction removed, the pellet was resuspended in 1 ml of 20% sucrose, 50 mM Tris pH 8 and sonicated with a Sonic Dismembrator Model 100 (Fisher Scientific) 3X for 5 sec at power setting 4. Samples were centrifuged 3X at 12,000 × g for 5 min at 4°C to remove unlysed cells. The supernatant was then centrifuged at 56K rpm for 1 h at 4°C in a Beckman TLA100.3 rotor. The supernatant containing the cytoplasmic fraction was removed and the pellet containing the membrane fraction was resuspended in 1 ml of 20% sucrose, 10mM Tris pH 8 by sonication. Equal volumes of fractions were assayed by immunoblotting with polyclonal *α*-AzuC antibody.

### Microscopy

Cells grown as indicated were harvested, resuspended in phosphate buffered saline (PBS) (KD Medical) and placed on lysine-coated glass bottom dish (Mattek Corporation). Cells were fixed by applying a 1% agarose pad on top of the sample with gentle pressure. Cells were viewed with a DeltaVision Core microscope system (Applied Precision) equipped with an environmental control chamber. Bright field and fluorescence images were captured with a Photometrics CoolSnap HQ2 camera. Seventeen planes were acquired every 0.2 μm at 22°C, and the data were deconvolved using SoftWorx software (GE Healthcare).

### Purification of chromosomally-encoded AzuC-SPA

Cells expressing AzuC-SPA (GSO351) cells grown in LB at 37°C overnight culture were diluted 1:100 into 1 l of M63 glucose minimal media and incubated at 37°C. At OD_600_ ∼1.0, cells were collected by centrifugation (4,650 × g, 20 min). The pellet was resuspended in 20 ml of TNG buffer [10 mM Tris (pH 7.5), 100 mM NaCl, 10% glycerol] supplemented with Protease Inhibitor Cocktail (Roche). The cells were lysed using a microfluidizer processor (Microfluidics) at 20,000 psi, and the insoluble cellular debris was removed by centrifugation (20,000 × g, 30 min). The cleared lysate was incubated with 50 mM dodecyl β-D-maltoside (DDM) at 4°C for 2 h. Next, the DDM-supplemented lysate was incubated with 500 µl of calmodulin-sepharose beads (Amersham Biosciences) overnight at 4°C. The lysate and beads were applied to a Bio-Spin disposable chromatography column (Bio-Rad Laboratories) and allowed to drain by gravity. The calmodulin column was washed 15 ml of TNG buffer with 2 mM DDM, 5 mM β-ME, and 2 mM CaCl. Finally, proteins were eluted from the calmodulin column in 1 ml TNG buffer supplemented with 4 mM EDTA, 5 mM β-ME, and 2.5% SDS. To analyze the protein samples, 7.5 μl of 2X Laemmli buffer was added to 21 μl of each sample. The samples were heated at 95°C for 5 min, and aliquots were subjected to SDS/PAGE in a 10–20% Tris-glycine gel (Invitrogen) at 12 V/cm. Proteins were visualized with Coomassie Blue Stain. Bands of interest were excised from the gel and analyzed by liquid chromatography-tandem mass spectrometry (LC-MS/MS). An identical purification was carried out for cells with chromosomal *acrZ-SPA* (GSO350) grown in 1 l of LB to OD_600_ ∼0.6.

### Purification of chromosomally-encoded GlpD-HA-His_6_ and MgtA-HA

MG1655 cells or cells expressing AzuC-SPA (GSO351), GlpD-HA-His_6_ (GSO1011) or the control MgtA-HA (GSO785) from the chromosome grown in LB at 37°C for 16 h, were diluted 1:100 into 1 l of M63 glucose minimal medium, M63 glycerol minimal medium or N medium with 500 µM MgSO_4_, respectively and incubated at 37°C. The WT strain and strains expressing GlpD-HA-His_6_ and AzuC-SPA were grown to OD_600_ ∼1.0. The strains expressing MgtA-HA were grown to OD_600_ ∼0.4–0.6, collected, washed 2X in N medium without added MgSO_4_, resuspended in N medium without added MgSO_4_ and grown for another 2.5 h to induce MgtA-HA expression. For all cultures, cells were collected by centrifugation (4,650 × g, 20 min) and resuspended in 15 ml of TNG buffer supplemented with Protease Inhibitor Cocktail (Roche). Cells from the SPA tagged protein cultures were mixed with the control WT or HA-tagged protein cultures at a 1:1 ratio. To ensure thorough mixing, cells were shaken gently at 4°C for 15 min. The cells were then homogenized as for the SPA-tagged protein purification and incubated with 50 mM DDM in 4°C for 2 h. The insoluble cellular debris was removed by centrifugation (20,000 × g, 20 min). Subsequently, the supernatant was applied to 100 µL of Pierce *α*-HA magnetic beads (Thermo Scientific) in a 50 ml tube and incubated overnight at 4°C. Beads were collected with a MagneSphere technology magnetic separation stand (Promega) and resuspended in 1 ml of TNG buffer. The beads were washed with 1 ml of TNG buffer (10X). The beads were then resuspended in 1XPBS (50 µl) and 2X Laemmli buffer (50 µl) and heated at 95°C for 5 min. Samples (15 µl) were analyzed on immunoblots using *α*-His or M2 *α*-FLAG antibodies.

### Dehydrogenase activity assay

Cells were grown in M63 glucose minimal medium to OD_600_ ∼1.0. Cells were pelleted and washed with M63 glycerol medium, pH 7.0 or pH 5.5. Cells were then resuspended in same volume of the same medium and grown at 37°C for 3 h. Cells (500 µl) were pelleted and resuspended in 500 µl of lysis buffer (25 mM Tris-HCl, 10 mM NaCl and 0.4% Triton X100). Cells were lysed by adding 0.6 g of glass beads and vortexing 30 s followed by 30 s incubation on ice, repeated 5X. The cells were then centrifuged at 20,000 × g for 2 min at 4°C, and the lysate was used to measure the dehydrogenase activity. A method monitoring MTT reduction to quantitate the dehydrogenase activity of GlpD (Yeh *et al*., 2008) was modified as follows. Each 225 µl microcuvette contained the following: 25 mM Tris/HCl pH 7.4, 100 mM NaCl, 1 mM MTT (Sigma Aldrich), 3 mM phenazine methosulfate (PMS, Sigma Aldrich) and 100 µl of lysate. This was used as the blank, and the reaction was initiated by the addition of 3.7 mM *sn*-glycerol-3-phosphate (Sigma Aldrich). The reduction of MTT at 570 nm was continuously monitored on a BMG LABTECH plate reader for 118 min at room temperature.

### β-galactosidase assays

Cultures were grown in LB to OD_600_ ∼1.0 with arabinose (0.2%). 100 µl of cells were added to 700 µl of Z buffer (60 mM Na_2_HPO_4_, 40 mM NaH_2_PO_4_, 10 mM KCl, 1 mM MgSO_4_, 50 mM β-mercaptoethanol). After the addition of 15 µl of freshly-prepared 0.1% SDS and 30 µl of chloroform, each sample was vortexed for 30 s and then incubated at room temperature for 15 min to lyse the cells. The assay was initiated by adding 100 µl of ONPG (4 mg/ml). The samples were incubated at room temperature until the reaction was terminated by the addition of 500 µl of 1M Na_2_CO_3_. A_420_ and A_550_ values determined with a spectrophotometer were used to calculate Miller units.

### Growth curves

Colonies of *ΔazuC::kan* (GSO193) transformed with pRI, pRI-AzuCR, pRI-AzuCR_L3STOP_, pKK, pKK-AzuC or pKK-AzuC_L3STOP_ grown on LB plates were inoculated into glucose (pH 7.0), glycerol (pH 7.0 and 5.5), and galactose (pH 7.0) and allowed to grow overnight at 37°C, at which point all cultures were in stationary phase. Cultures were diluted to OD_600_ ∼0.05 (time 0) in 25 ml of the same media and grown at 37°C. OD_600_ was measured at 16 h or growth was followed for 29 h.

### GFP reporter assay

The GFP reporter assay was principally done as described previously (Corcoran *et al*., 2012, Urban & Vogel, 2009). WT or *ΔazuC::kan* (GSO193) cells were transformed with a *cadA-gfp*, *cadA-gfp-M1*, *galE-gfp* or *galE-gfp-M2* reporter plasmid and a pRI-AzuCR, pRI-AzuCR_L3STOP_, pRI-AzuCR_L3STOP_-M1 or AzuCR_L3STOP_-M2 over-expressing plasmid or pRI as a control. Single colonies were grown overnight at 37°C in LB supplemented with ampicillin and chloramphenicol. The cultures were diluted to OD_600_ ∼0.05 in fresh medium and grown at 37°C for 3 h in a 96 deep-well plate. An aliquot (1 ml) of each culture was centrifuged and the pellet was resuspended in 220 µl of 1X PBS. Fluorescence was measured using the CytoFLEX Flow Cytometer (Beckman Coulter). Three biological repeats were analyzed for every sample.

### Hfq and ProQ co-immunoprecipitation assays

Cell extracts were prepared from MG1655 cells grown in M63 glucose medium to OD_600_ ∼0.5. Cells corresponding to the equivalent of 20 OD_600_ were collected, and cell lysates were prepared by vortexing with 212-300 μm glass beads (Sigma-Aldrich) in a final volume of 1 ml lysis buffer (20 mM Tris-HC, pH 8.0, 150 mM KCl, 1 mM MgCl_2_, 1 mM DTT). Immunoprecipitations were carried according to (Zhang *et al*., 2002) using 100 µl of Hfq antiserum (Zhang *et al*., 2002) or 100 µl of ProQ antiserum (Melamed *et al*., 2020), 120 mg of protein-A-sepharose (Amersham Biosciences, Piscataway, NJ) and 950 µl of cell extract per immunoprecipitation reaction. Immunoprecipitated RNA was isolated from immunoprecipitated pellets by extraction with phenol:chloroform:isoamyl alcohol (25:24:1, pH 8), followed by ethanol precipitation. Total RNA was isolated from 50 μl of cell lysate by Trizol (Thermo Fisher Scientific) extraction followed by chloroform extraction and isopropanol precipitation. Total and co-IP RNA samples were resuspended in 20 μl of DEPC H_2_O and 2 µg of total RNA or 200 ng of IP RNA was subjected to Northern analysis as described below.

## Acknowledgements

We thank P. Backlund for carrying out mass spectrometry in the *Eunice Kennedy Shriver* National Institute of Child Health and Human Development core, A. Zhang for assistance with the co-immunoprecipitation assays, T. Updegrove for assistance with microscopy, A. Buskirk for the *azuC* browser image, and M. Hemm, N. Soltanzad, L. Stamper and A. Romero for help with initial experiments, and members of the Storz group for comments on the manuscript. This research was supported by the Intramural Research Program of the *Eunice Kennedy Shriver* National Institute of Child Health and Human Development.

## Author contributions

Studies were planned by all authors and carried out by MR, JA, SB and BJP. MR, JA and GS prepared the manuscript.

## Conflict of interest

The authors declare that they have no conflict of interest.

## Expanded View Figures

**Figure EV1.**
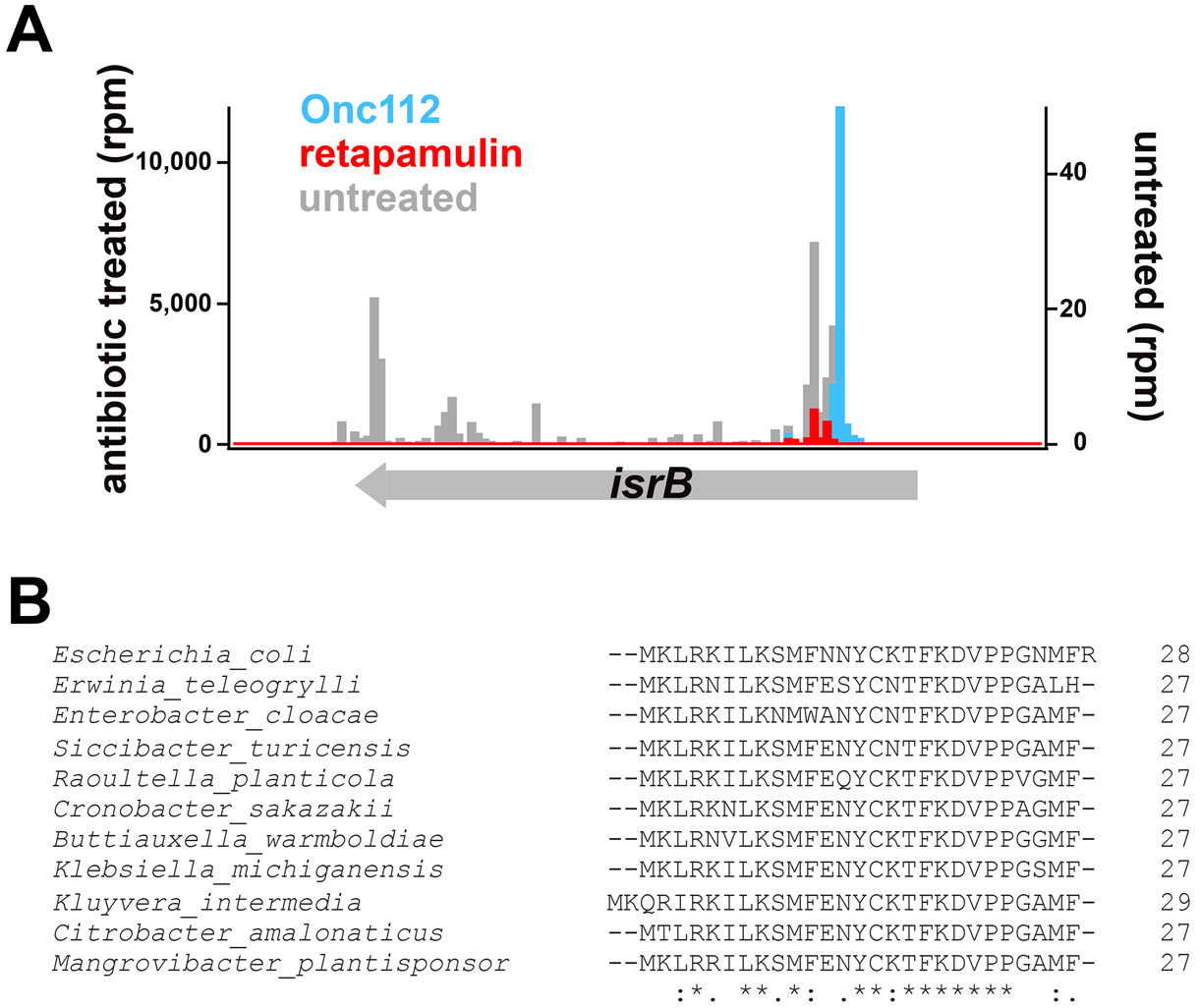
Ribosome binding to and conservation of *azuC*. A The *azuC* open reading frame lies within a region that was previously reported to encode the IsrB sRNA (Chen *et al*., 2002). Translation is detected by ribosome density on the *isrB* gene for an untreated control (gray) (Weaver *et al*., 2019), and cells treated with the translation inhibitors Onc112-treated (blue) (Weaver *et al*., 2019) or retapamulin-treated (red) (Meydan *et al*., 2019). B The AzuC amino acid sequences from *E. coli* K12 and other bacterial species aligned with ClustalW (Madeira *et al*., 2019). “*” indicates the residues are identical in all sequences and “:” and “.” respectively indicate that conserved and semi-conserved substitutions as defined by ClustalW.

**Figure EV2.**
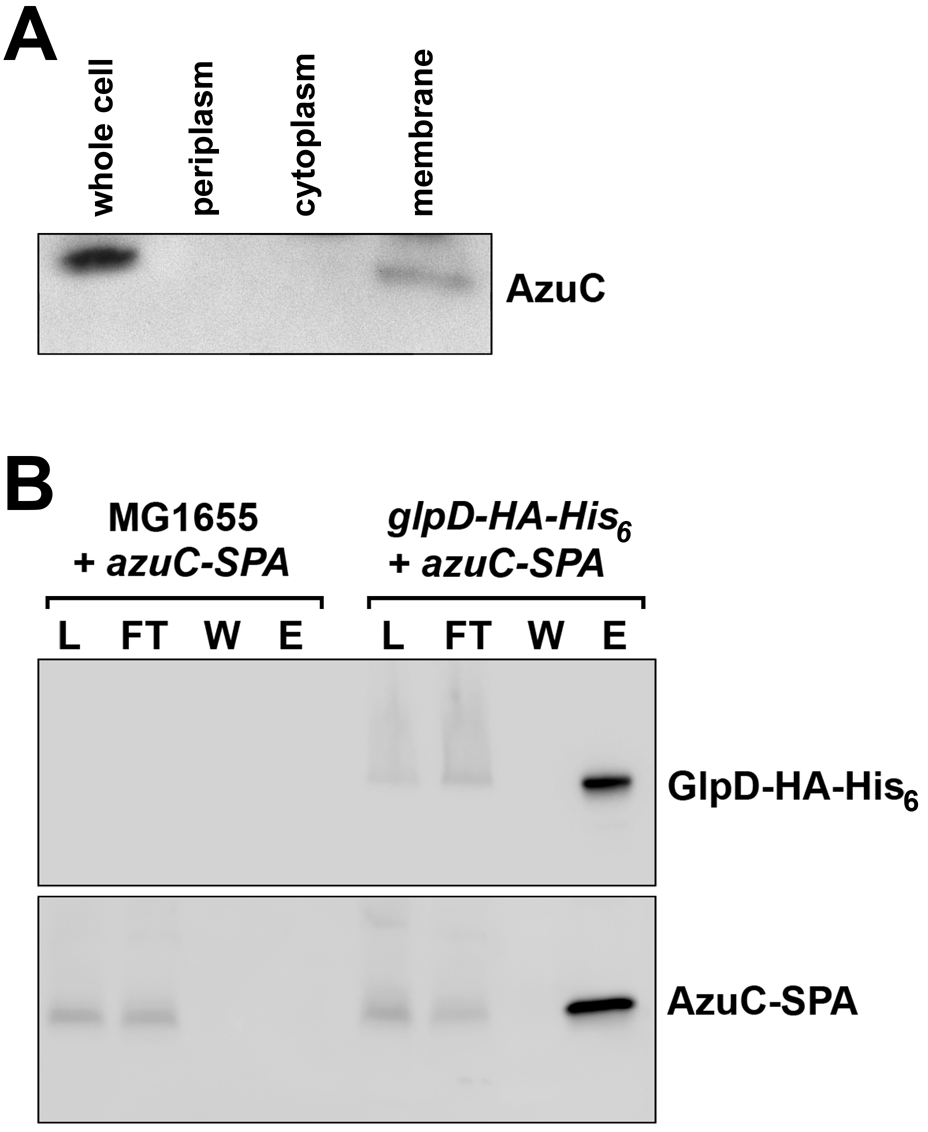
Fractionation showing subcellular localization of untagged AzuC and AzuC-SPA co-purification with GlpD-HA-His_6_ compared to untagged control strain. A AzuC was overexpressed at low levels from the arabinose-inducible P_BAD_ promoter on the multicopy pAZ3 plasmid derivative of pBAD18 (Kawano *et al*., 2005). After induction with arabinose, cell extracts were fractionated into periplasmic, cytoplasmic, and membrane fractions. The fractions were then examined on immunoblots using polyclonal *α*-AzuC primary antibody followed by *α*-rabbit secondary antibody. AzuC expressed from the chromosome could not be detected by the polyclonal *α*-AzuC antibody. B AzuC-SPA cells grown in M63 glucose and MG1655 cells or GlpD-HA-His_6_ cells grown in M63 glycerol, to OD_600_ ∼1.0 were mixed in a 1:1 ratio. The mixed cells were homogenized, cell lysates (L) were applied to anti-HA beads and the flow-through (FT) fractions were collected. The beads were washed (W), after which the bound proteins were eluted (E) and examined on immunoblots using either *α*-HA antibodies to detect GlpD-HA-His_6_ (top panel) or *α*-FLAG antibodies to detect AzuC (bottom panel).

**Figure EV3.**
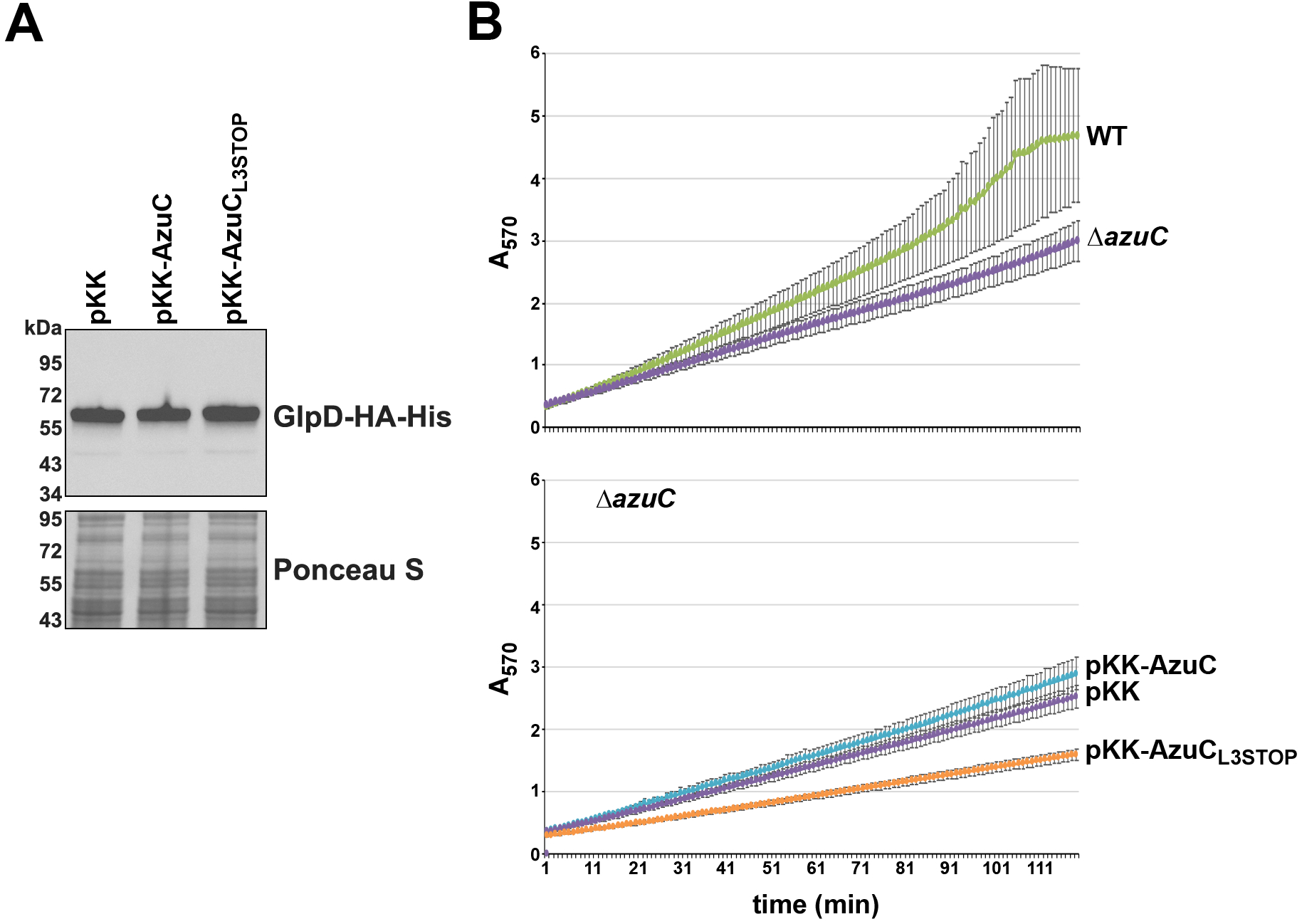
Effect of AzuC overexpression on GlpD levels at pH 5.5 and on GlpD activity at pH 7.0. A *ΔazuC* cells expressing GlpD-HA-His_6_ (GSO1013) were transformed with pKK, pKK-AzuC, and pKK-AzuC_L3STOP_, grown in M63 glycerol medium to OD_600_ ∼0.5 and examined on immunoblots using *α*-HA antibodies to detect GlpD-HA-His_6_. The membrane was stained with Ponceau S stain to control for loading. B MG1655 or *ΔazuC* (GSO193) cells (top panel) or *ΔazuC* (GSO193) cells transformed with pKK, pKK-AzuC, or pKK-AzuC_L3STOP_ (bottom panel) were grown in M63 glucose medium to OD_600_ ∼0.5 and then washed and resuspended in M63 glycerol medium at pH 7.0. Cells were allowed to grow for an additional 3 h at which point the dehydrogenase activity assay was performed.

**Figure EV4.**
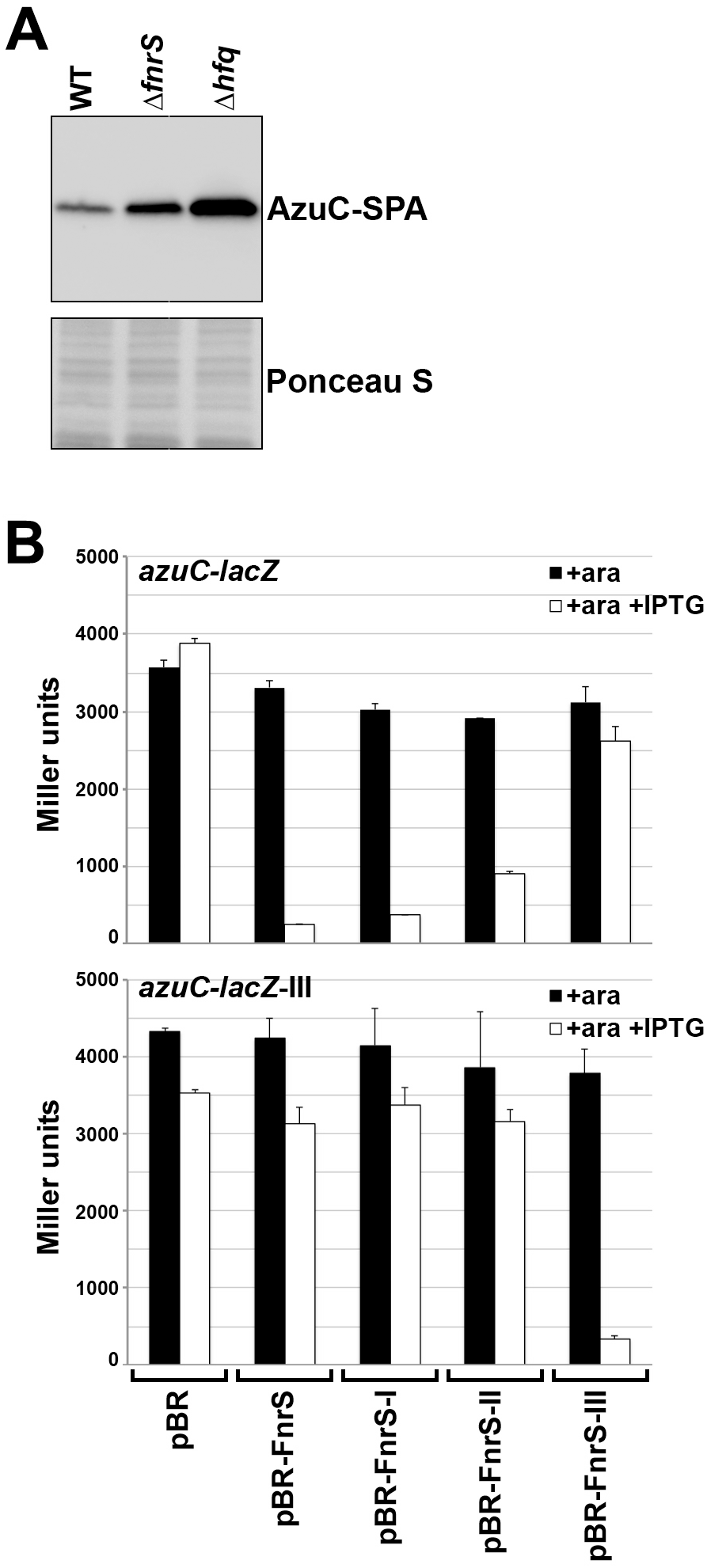
FnrS sRNA represses AzuC expression. A AzuC-SPA levels in strains lacking Hfq or FnrS. AzuC-SPA levels in WT (GSO351), *Δhfq* (GSO1007) or Δ*fnrS* (GSO1023) strains grown in LB to OD_600_ ∼0.5. *α*-FLAG antibody was used to detect the SPA tag. B β-galactosidase activity was assayed for P_BAD_*-*5’-UTR*_azuC_*-*lacZ* (GSO1024) or P_BAD_*-*5’-UTR*_azuC_*-*lacZ* -III (GSO1025) cells carrying pBR, pBR-FnrS or pBR-FnrS mutants (pBR-FnrS I, II or III). Cells were grown to OD_600_ ∼0.4-0.5 and treated with either just arabinose (0.2%) (black bars) or arabinose (0.2%) and IPTG (1 mM) (white bars), and cells were grown another 40 min. The average of three independent trials is shown, and the error bars represent one SD.

**Figure EV5.**
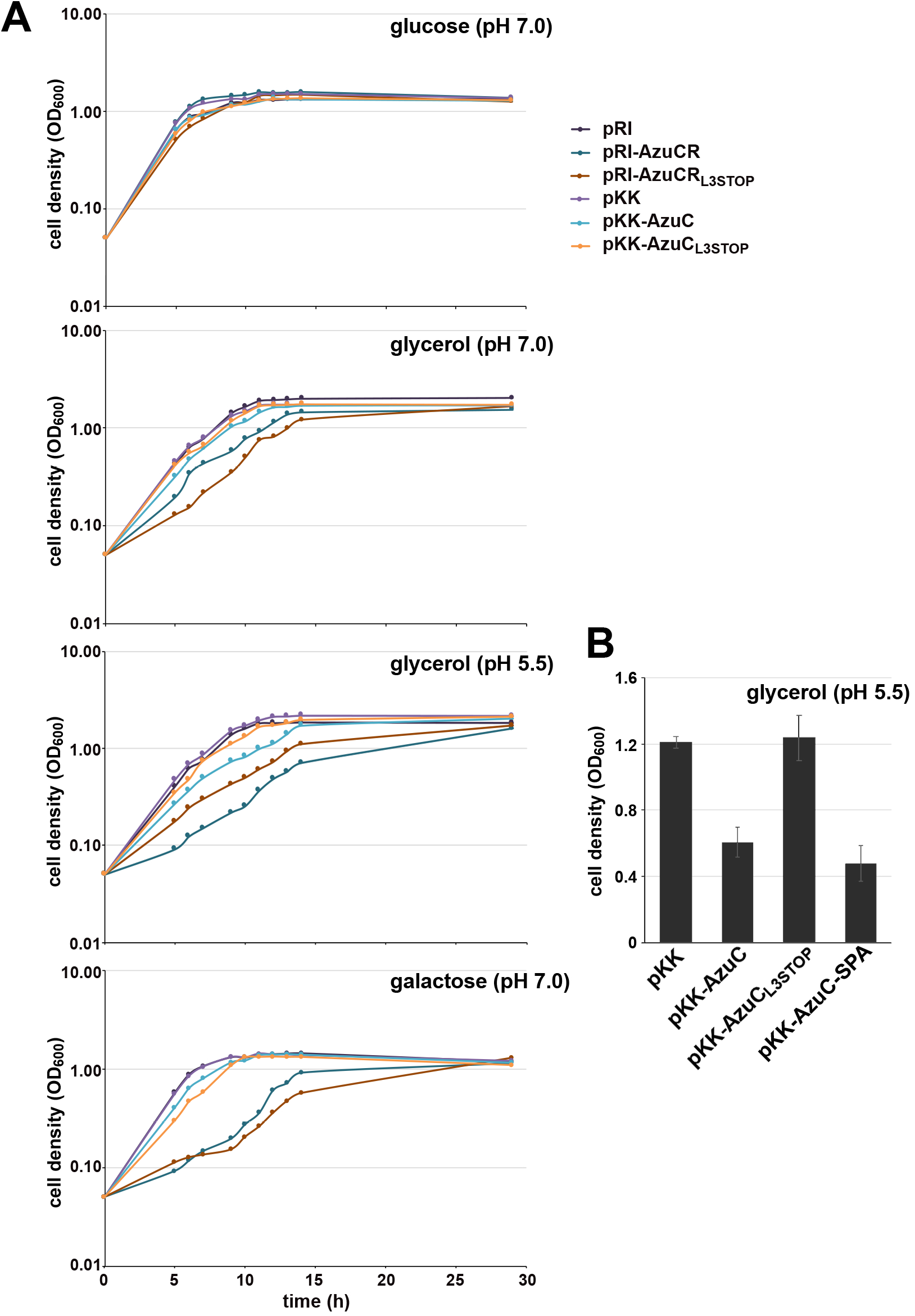
Growth curves for AzuC and AzuCR overexpression. A Δ*azuC::kan* strain (GSO193) transformed with pRI, pRI-AzuCR, pRI-AzuCR_L3STOP_, pKK, pKK-AzuC, or pKK-AzuC_L3STOP_ was grown in M63 medium with different carbon sources: glucose (pH 7.0), glycerol (pH 7.0 and 5.5), and galactose (pH 7.0) and growth was tracked by OD_600_ over 30 h. B Growth of the *ΔazuC::kan* strain (GSO193) transformed with pKK, pKK-AzuC, pKK-AzuC_L3STOP_, or pKK-AzuC-SPA in M63 medium with glycerol pH 5.5 was measured at 16 h after dilution by OD_600_.

## Supplemental Tables

Appendix Figure S1. Strains used in study.

Appendix Figure S2. Plasmids used in study.

Appendix Figure S3. Oligonucleotides used in study.

